# Autogenous and anautogenous *Culex pipiens* bioforms exhibit insulin-like peptide signaling pathway gene expression differences that are not dependent upon larval nutrition

**DOI:** 10.1101/2021.06.20.449144

**Authors:** Alys M. Cheatle Jarvela, Katherine L. Bell, Anna Noreuil, Megan L. Fritz

## Abstract

*Culex pipiens* form *pipiens* and *Cx. pipiens* form *molestus* differ in their ability to produce eggs without a bloodmeal. Autogenous mosquitoes, such as the *molestus* bioform of *Cx. pipiens*, depend on nutrition acquired as larvae instead of a bloodmeal to fuel the energy intensive process of vitellogenesis, which requires abundant production of yolk proteins. In anautogenous mosquito systems, ovary ecdysteroidogenic hormone (OEH) and insulin-like peptides (ILPs) transduce nutritional signals and trigger egg maturation in response to a bloodmeal. It is unclear to what extent the process is conserved in autogenous mosquitoes and how the bloodmeal trigger has been replaced by teneral reserves. Here, we measured the effects of a series of nutritional regimens on autogeny, time to pupation, and survival in *Cx. pipiens* form *molestus* and form *pipiens*. We find that abundant nutrients never result in autogenous form *pipiens* and extremely poor food availability rarely eliminates autogeny from form *molestus*. However, the number of autogenous eggs generated increases with nutrient availability. Similarly, using qPCR to quantify gene expression, we find several differences in the expression levels of *ilps* between bioforms that are reduced and delayed by poor nutrition, but not extinguished. Changes in *OEH* expression do not explain bioform-specific differences in autogeny. Surprisingly, the source of most of the gene expression differences correlated with autogeny is the abdomen, not the brain. Overall, our results suggest that autogeny is modulated by nutritional availability, but the trait is encoded by genetic differences between forms and these impact the expression of ILPs.

## Introduction

Blood is a requirement for egg production in most mosquito vectors of disease, and acquisition of blood drives pathogen transmission. This reproductive strategy is known as anautogeny. Yet egg production in the absence of a blood meal, or autogeny, has been described in dozens of mosquito species (Vinogradova, 2000). Autogeny can be obligatory, where females always lay their first (and, in some species, subsequent) batches of eggs without a blood meal, or facultative, where females may lay an autogenous egg raft dependent on their larval environmental conditions (Attardo et al., 2005; Provost-Javier et al., 2010; Tsuji et al., 1990). While it is well-established that autogenous reproduction is under genetic control (Aslamkhan and Laven, 1970; Krishnamurthy and Laven, 1961; Mori et al., 2008; O’Meara et al., 1969; Spielman, 1957; Trpis, 1978), the molecular basis for this unique reproductive strategy remains unknown. Comparing the molecular and physiological tradeoffs underlying autogenous and anautogenous reproduction has strong potential to elucidate the key genetic changes that led to multiple instances of reproductive divergence within and among mosquito species (O’Meara, 1985).

Due to its epidemiological relevance, the reproductive physiology of anautogenous mosquitoes has been investigated extensively. Egg development requires that yolk proteins produced by the fat body are deposited into developing oocytes, a process known as vitellogenesis. In anautogenous mosquitoes, vitellogenesis is tightly suppressed until a blood meal is obtained (reviewed in Attardo et al., 2005; Hansen et al., 2014). Four key signaling pathways, juvenile hormone III (JH), ecdysone, insulin-signaling, and target of rapamycin (TOR), interact to regulate the expression of yolk protein precursors (YPPs) in the fat body, vitellogenesis, and ovarian maturation (reviewed in Attardo et al., 2005; Hansen et al., 2014). JH acts on the fat body and ovaries during maturation, influencing the ability of a female to activate *YPP* genes (Noriega, 2004; Zou et al., 2013). Once a female obtains a blood meal, the steroid hormone ecdysone is the major hormone that regulates egg maturation via activation of target genes, such as the predominant *YPP* gene, *vitellogenin* (*Vg*) (reviewed in Roy et al., 2016). Ecdysone is regulated synergistically by ovary ecdysteroidogenic hormone (OEH) and insulin-signaling. After feeding, stretch receptors in the gut trigger the release of OEH from the brain into the hemolymph, which stimulates the production and release of ecdysone by follicle cells of the ovary (Dhara et al., 2013; Hagedorn et al., 1979, 1975). Insulin-like peptides (ILPs), a group of evolutionarily conserved peptide hormones, may also influence *YPP* gene expression both directly and indirectly (Hansen et al., 2014).Together, insulin pathway and ecdysone signaling activate *YPP* expression in the fat body (Roy et al., 2007). In *Aedes aegypti*, one member of the insulin-like peptide family, ILP3, has been demonstrated to be essential for the production of ecdysone, providing indirect reinforcement of *YPP* expression (Brown et al., 2008, Dhara et al. 2013, Vogel et al. 2015). ILP3 also stimulates serine protease activity in the midgut for blood meal digestion, contributing to the availability of amino acids required to produce YPPs (Gulia-Nuss et al., 2012). The TOR signaling pathway detects these increases in amino acid levels and contributes to the activation of *YPP* gene expression (Hansen et al., 2004). Ultimately, vitellogenesis is not initiated in anautogenous females until these nutrient sensing pathways signal the availability of yolk building blocks after a blood meal.

In contrast to anautogenous reproduction, the physiology of ovarian maturation in autogenous mosquitoes has received less attention (Gulia-Nuss et al., 2015, 2012; Kassim et al., 2012; Provost-Javier et al., 2010). One of the most well-studied autogenous species is the rockpool mosquito, *Georgecraigius atropalpus* (formerly *Aedes* and *Ochlerotatus)(Gulia-Nuss et* al., 2015, 2012; Telang et al., 2013, 2006). *G. atropalpus* females that are decapitated within 6 hours of pupal emergence fail to initiate vitellogenesis, demonstrating that hormones secreted from the brain are critical to autogenous reproduction in this species. When injected, OEH fully restores and ILP3 partially restores reproductive maturation in these decapitated females (Gulia-Nuss et al., 2012), demonstrating that the roles of these hormones in *G. atropalpus* are analogous to what has been found in the primarily anautogenous *Ae. aegypti* after they have acquired a blood meal.

In *G. atropalpus* and other autogenous mosquitoes, nutritional status interacts with genetic factors, including those underlying hormonal secretion from the brain, to influence autogenous egg production. Females emerging from crowded, or nutrient poor larval habitats may fail to reproduce autogenously or have low fecundity (Kassim et al., 2012; Krishnamurthy and Laven, 1961; Lounibos et al., 1982; O’Meara and Krasnick, 1970; Trpis, 1978). Such nutritional control of trait expression is observed in other insect species (Casasa and Moczek, 2018; Chandra et al., 2018; Wheeler and Frederik Nijhout, 1983), and in some cases, the molecular mechanism underlying this nutritional regulation is known. For example, horn size in taurus scarab beetles (*Onthophagus taurus*) is dependent on nutritional state, which is transduced by insulin signaling (Casasa and Moczek, 2018). In ants, upregulation of *insulin-like peptide 2* (*ilp2*) occurs in reproductive as opposed to non-reproductive individuals and larval nutrition can influence adult ILP2 expression levels (Chandra et al., 2018). In these and other examples, an emerging theme is that insulin signaling is a critical component of the mechanism that ties phenotype to nutritional status (reviewed in Nijhout and McKenna, 2018). In autogenous mosquitoes, it is unclear how and whether the nutritional state of females impacts the expression of ILPs and other genes underlying induction of vitellogenesis and ovarian maturation.

Here, we examine the interaction between nutrition, gene expression, and ovarian maturation in a species of mosquito which is polymorphic for autogeny, *Culex pipiens*. Within the species, there are two interfertile and morphologically indistinguishable bioforms that exhibit divergent reproductive strategies (*Culex pipiens* form *pipiens* and form *molestus*, hereafter *pipiens* and *molestus*) (Harbach et al., 1984; Spielman, 2001; Spielman and Wong, 1973). The form *molestus* is thought to be facultatively autogenous, while the form *pipiens* is anautogenous (Roubaud, 1929). Although these populations are genetically distinct (Fonseca et al., 2004; Kent et al., 2007; Yurchenko et al., 2020), the molecular mechanisms underlying their divergent reproductive strategies have not been identified.

We begin to address this knowledge gap by quantifying changes in expression of key nutrient-sensing pathway genes for the autogenous *molestus*, as compared to the anautogenous *pipiens*. Multiple genes previously demonstrated to play a critical role in ovarian maturation, including *oeh*, the full suite of *ilps*, their downstream effector *foxo*, as well as *Vg1b*, were examined for expression level differences in female heads and abdomens for four days following adult emergence. We also measured gene expression in females that experienced nutrient rich and poor larval conditions, which produced dramatic changes in time spent as larvae and fecundity in autogenous *molestus*. Expression patterns for several of these genes differed temporally, by form, and in a body segment specific way. Interestingly, insulin-signaling pathway genes whose expression differed by form also show delayed or dampened expression in poorly-fed *molestus* individuals, but never matched the expression levels observed in well-fed *pipiens*. This demonstrates the relatively greater importance of genotype over nutrition in autogenous egg production for *Cx. pipiens*. Furthermore, this work establishes a system that will allow investigation of the key genetic variants which halt ovarian maturation in some females yet facilitate it in others. Such variants would make attractive targets for novel genetic control measures that limit ovarian maturation, egg production, and ultimately vector population growth.

## Results and Discussion

### *Bioinformatic and phylogenetic analyses reveal six* Culex ilp *orthologs*

At the start of this work, three *ilps* had been identified in *Cx. pipiens* (named *ilp1, ilp2*, and *ilp5*)(Sim and Denlinger, 2009), but based on the numbers in *Aedes aegypti* (n = 8; (Riehle et al., 2006)), *Anopheles gambiae* (n = 5; (Riehle et al., 2002)), *Anopheles stephensi* (n = 5; (Marquez et al., 2011)), and *Drosophila melanogaster* (n = 8; Brogiolo et al., 2001; Colombani et al., 2012; Garelli et al., 2012; Grönke et al., 2010), it was predicted that more remained to be discovered (Sharma et al., 2019; Sim and Denlinger, 2009). We identified six *Culex ilp* genes by performing a HMMR analysis in Vectorbase using an alignment of all eight *Drosophila* and *Aedes ilps* against the CpipJ2.4 geneset. A phylogenetic analysis of these six *Culex* ILPs determined their orthology to established sequences from *Drosophila, Aedes*, and *Anopheles* (Fig. 1). We confirmed the identity of previously isolated *Cpip-ilp1* and *-ilp5* genes, as well as those from the *Culex quinquefasciatus* Johannesburg genome that were included in other’s phylogenetic analyses (Marquez et al., 2011; Sim and Denlinger, 2009). One gene previously named *Cpip-ilp2* (Genbank accession ACM66967.1) was orthologous to *ilp3* in other mosquito species. Comparison of *“Cpip-ilp2”* to *Cqui-ilp2* and *Cqui-ilp3* further supports its orthology with *ilp3* (Fig. S1). Therefore, we refer to this sequence as *Cpip-ilp3* in this work.

**Figure 1:**
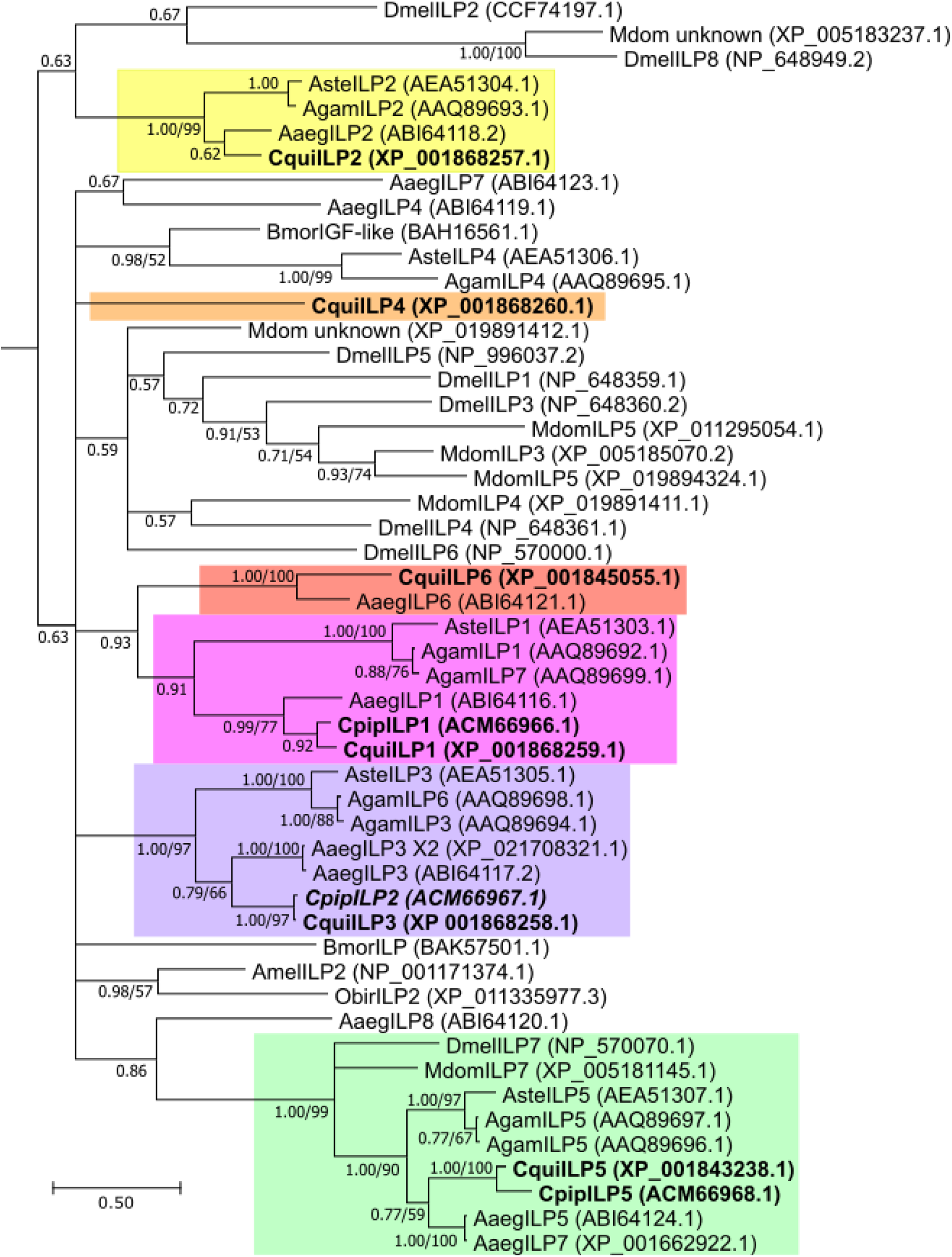
Phylogenetic Tree of Dipteran ILPs establishes *Culex ilp* orthology. Six distinct *Culex ilp* genes (bold) were assigned orthology based on clustering with their closest homologs in other species, indicated by colored boxes around each clade. *Cqui-ilp4* is a lineage-specific gene with no homology to other Dipteran *ilp* genes (orange box). Only *Cqui-ilp5* has well supported orthology to a *Dmel-ilp* gene (*ilp7*) (green box). Tree topology determined by TOPALI v2.5 using a MrBayes algorithm. Posterior probabilities listed at nodes to indicate statistical support. When the node was also supported by a maximum likelihood tree (>50) (TOPALI, PhyML algorithm, JTT + G, 100 bootstrap runs), the bootstrap support is listed as the second value at the node. Genbank accession numbers listed next to gene names.

One new *ilp* gene was identified and had likely been overlooked in previous work because it is unique to *Culex* (XP_001868260.1). We named this gene *ilp4* because it is syntenic with *ilp*s *1-3*, as is the case in other Dipteran genomes (Fig. S2). It is not a clear ortholog of any other Dipteran *ilp4*, however, and likely arose by a lineage-specific gene duplication event. All of the genes in this syntenic cluster have sequence features that define them as “insulin-like peptides”, such as a long, cleavable C peptide (Okamoto and Yamanaka, 2015). *Cpip-ilp5* and other mosquito *ilp5* orthologs are not syntenic in the genome with other *ilp*s and contain distinct sequence features that define them as orthologs of *Dmel-ilp7* (Krieger et al., 2004; Okamoto and Yamanaka, 2015; Riehle et al., 2006). Our phylogenetic analysis upholds this grouping (Fig. 1). Similarly, *Cqui-ilp6* is in a clade with *Aaeg-ilp6* in our analysis and both are located elsewhere in the genome, not in a syntenic cluster with other ILP genes. This is also true for *Dmel-ilp6*, which is considered “IGF-like” based on sequence features such as its short C peptide. Although our phylogeny does not group mosquito *ilp6* genes with *Dmel-ilp6*, the short C peptides, extended C-terminus following the A domain, and genomic locations suggest that they can also be considered “IGF-like” genes (Riehle et al., 2006). In sum, there are six *ilp* genes in *Cx. pipiens* assemblage species: *ilps1-4* are true insulin-like peptides, *ilp5* is similar to *Drosophila’s* atypical *ilp7*, and *ilp6* is best described as an IGF-like peptide.

### Autogenic ovarian maturation is enhanced by a nutrient rich larval environment

Prior to quantifying gene expression, we confirmed that the timing and degree of *Culex* ovarian maturation was consistent with previous observations of autogenous and anautogenous mosquito populations. Dissections of adult *molestus* and *pipiens* females conducted over 96h post emergence (PE) showed that ovarian maturation in *molestus* progressed beyond that of *pipiens* by 48h PE and was complete by 96h PE, regardless of mating status (Fig. 2). Failure of mating status to impact autogenous ovarian maturation agreed well with previous studies of another *molestus* population and *G. atropalpus* (Gulia-Nuss et al., 2012; Kassim et al., 2012; Spielman, 1957). Furthermore, our time course of ovarian maturation was consistent with studies showing *molestus* follicle length reaches its maximum between 80-105h post-emergence (Spielman, 1957), and egg deposition begins at 120h (Kassim et al., 2012).

**Figure 2:**
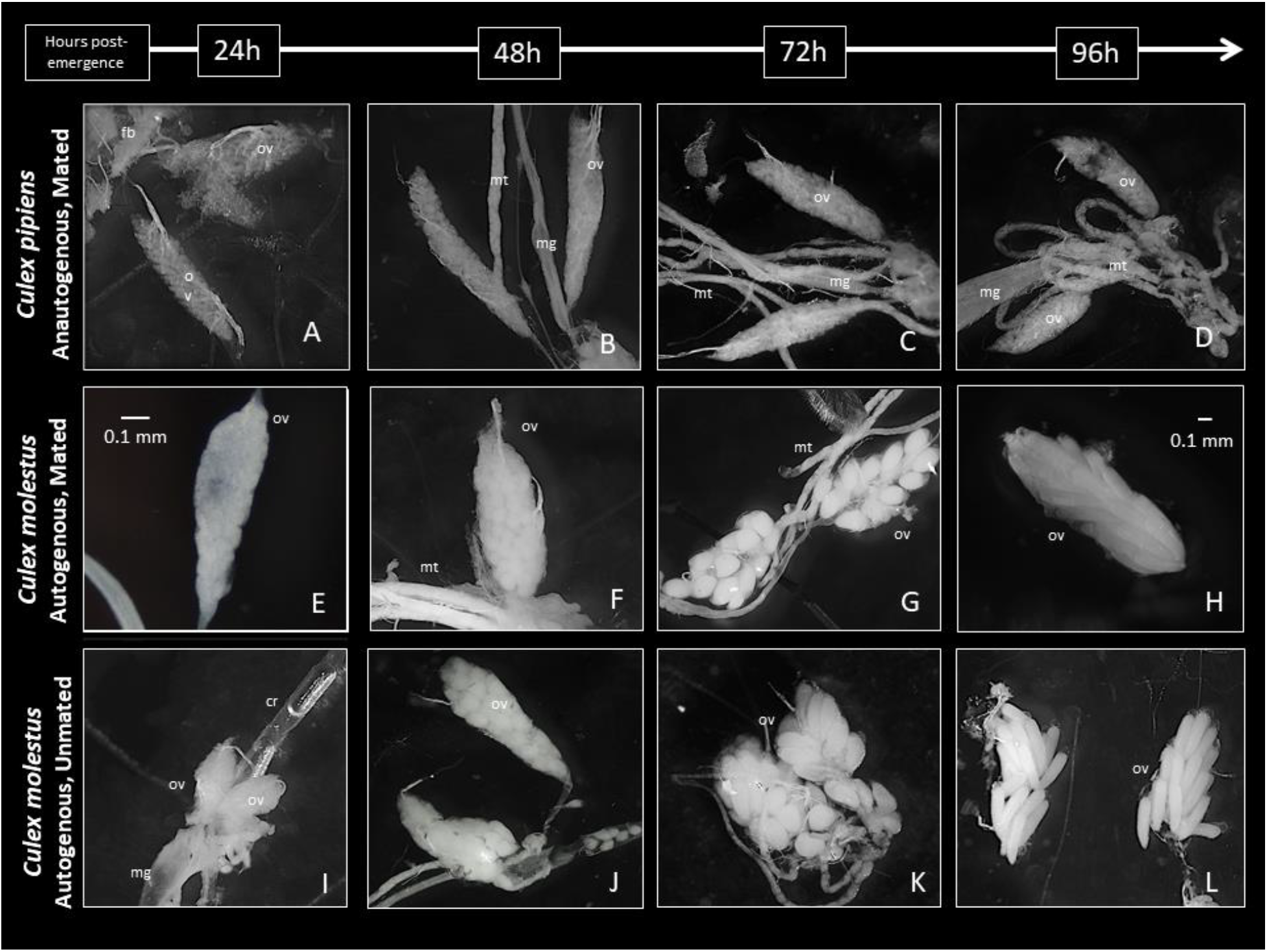
Ovarian development up to 96 hours post adult emergence in anautogenous *pipiens* and autogenous *molestus* females. Panels A-D depict ovarian development in anautogenous, form *pipiens* females in which follicles remain in resting stage (Christopher’s Stage - IIb) until a blood meal is taken. Panels E-H depict autogenous ovarian development for form *molestus* females provided the opportunity to mate. Panels I-L show ovarian development in autogenous *molestus* females denied a mating opportunity. A 0.1 mm scale is shown in panels E and H for reference. ov, ovary; mg, midgut; fb, fatbody; mt, malpighian tubules; cr, crop.

Nutrient rich and poor aquatic environments are known to impact body size, larval development, mortality, and egg production in autogenous mosquitoes (Kassim et al., 2012; Lounibos et al., 1982; O’Meara and Krasnick, 1970; Telang and Wells, 2004). Here, we examined both anautogenous *pipiens* and autogenous *molestus* for these traits in a comparative framework under identical environmental conditions. Rather than rearing larvae together in the same environment, as was previously done (Kassim et al., 2012; Spielman, 1957), we reared each larva in a single well of cell culture plate to avoid the confounding effects of competition on trait expression. Each larva was consistently fed one of four diet treatments ranging from 0.25mg desiccated liver powder (LP) + 0.14mg dry yeast (DY) per larva (extra-low) to 1.07mg LP + 0.53mg DY (high) every other day. For both forms, larval survivorship was high (> 85%) and did not differ by diet treatment (Fig. S3; Tables S1 & S2). Wing length, which serves as a proxy measure of female body size and teneral reserves (Telang et al., 2006) was positively correlated with larval nutrient availability, however (Table 1; Tables S3 & S4). Together, this indicated that our diet treatments effectively manipulated larval nutrition without inducing mortality.

**Table 1:** Mean wing length is positively correlated with nutrient availability in larval *Cx. pipiens*. Means and standard deviations were calculated using one wing for 12 individuals per form, per diet treatment. Correlations between wing length and larval nutrition were determined using a Bayesian generalized linear model (Tables S3 & S4).

| Form | Diet Treatment | Mean Wing Length (mm) | Standard Deviation |
| --- | --- | --- | --- |
| *Cx. pipiens* form *molestus* | High | 3.43 | 0.21 |
|  | Medium | 3.22 | 0.10 |
|  | Low | 3.06 | 0.11 |
|  | Extra Low | 2.84 | 0.09 |
| *Cx. pipiens* form *pipiens* | High | 3.66 | 0.15 |
|  | Medium | 3.51 | 0.16 |
|  | Low | 3.32 | 0.15 |
|  | Extra Low | 3.16 | 0.14 |

As expected based on previous work (Attardo et al., 2005), larval development time was negatively correlated with nutrient availability for both forms (Fig. S4; Tables S5 & S6). When fed the highest diet treatment, form *pipiens* and *molestus* larvae developed at similar rates (Tables S5 & S7), where mean days to pupation were 7.2 (s.d. = 0.8, n = 208) and 7.8 (s.d. 1.1, n = 222), respectively. At the lowest diet treatments, larval development times lengthened to 10.2 (s.d. = 1.9, n = 202) and 13.1 (s.d. 2.6, n = 200) days, respectively, and there was higher variation in development time for *molestus*. We reasoned that sex-specific differences in nutrient acquisition, particularly for *molestus*, could account for this additional variation. In a separate experiment, we quantified male and female development times for both *molestus* and *pipiens*, and our results revealed a three-way interaction between sex, form, and diet treatment (Fig. 3; Table S8 & S9). Female *molestus* larvae compensated for very low nutrient availability by lengthening their development time beyond what we observed for *pipiens* females and males of both forms (Fig. 3). This was consistent with previous studies of autogenous mosquitoes (Lounibos et al. 1982, Kassim et al. 2012), but unlike our results, *molestus* larvae from previous studies suffered increased mortality (up to 38.7%), likely due to the more severe starvation conditions imposed for the lowest diet treatments (Kassim et al., 2012).

**Figure 3:**
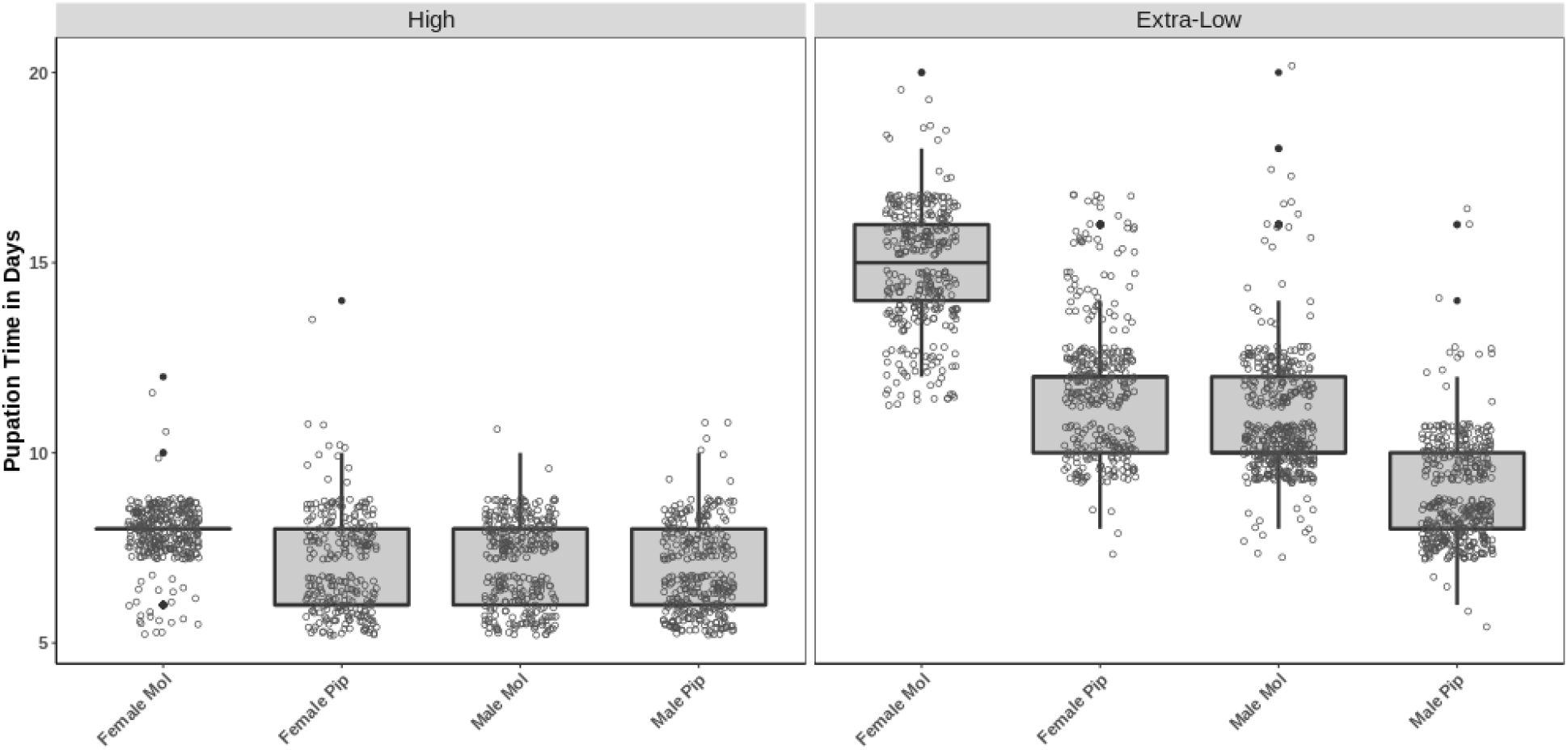
Days until pupation, per treatment, for *molestus* and *pipiens*. Males and females are shown separately. Unfilled points represent observations for individual larvae, while filled points represent outlier observations (1.5X greater or less than the interquartile range).

Ovarian dissections revealed that larval nutrition was positively correlated with the numbers of elongated follicles produced by each *molestus* female (Tables S10 & S11), while *pipiens* females never showed signs of ovarian maturation (Table S12). Form *molestus* females raised at the highest diet treatment matured, on average, 47.6 (s.d. = 14.3, n = 62) follicles whereas those from the lowest diet treatment matured only 14.3 (s.d. = 9.6, n = 51; Table S12, Fig. S4). We also quantified the probability that *molestus* females failed to produce any mature follicles according to diet treatment. From the highest to lowest diet treatment, 1.6 %, 3.2%, 2.8% and 19.6% of *molestus* females showed no signs of ovarian maturation. Our statistical analysis indicated that the lowest diet treatment had greater potential to halt ovarian maturation in *molestus* females than did the other diet treatments (Tables S13 & S14), confirming that expression of autogeny is the result of a genotype by environment interaction. Finally, our observation that poorly nourished *molestus* females lengthened larval development time led us to quantify the relationship between time spent as larvae and the numbers of elongated follicles produced. We tested for a positive correlation between development time and numbers of elongated follicles produced by females reared under the low and extra low diet treatments but found little evidence of this (Fig. S5; Tables S15 & S16).

Altogether, these results support and add to previous work, showing that autogenic follicle development in *Culex* is a form-specific trait that is quantitatively impacted and sometimes eliminated by larval diet (Fig. S4). Poor nutritional conditions lengthen larval development in a pronounced, sex-specific way for autogenous female *molestus*, compared to anautogenous *pipiens* reared under identical conditions (Fig. 3; Table S8). Yet increased larval development time does not necessarily result in strong increases in the numbers of elongated follicles produced by poorly nourished females. These results suggest there is a threshold level of nutrient acquisition that must be met prior to pupation in autogenous *molestus* females. In our hands, when nutritional requirements were met and female *molestus* pupated, most initiated follicle maturation upon eclosion. Extended larval development times observed for *molestus* females appeared to be more important for progression from larva to pupa than for increasing reproductive output, however (Fig. S5).

### Gene expression changes associated with autogeny are frequently observed in abdominal tissue samples and are damped or delayed under poor nutritional conditions

After observing significant, yet non-lethal impacts of diet on larval development and reproduction in our *Culex* forms, we compared the expression patterns of nine genes involved in nutrient-sensing and ovarian maturation for females raised under nutrient rich and poor conditions using our previous experimental design. These gene candidates included *ilps 1-6, foxo, Vg1b*, and *oeh*. Autogenous *molestus* and anautogenous *pipiens* were reared individually under the highest or lowest diet treatments as described above, and non-bloodfed females were collected at 24-hour intervals PE in groups of ten, according to form and diet treatment. Heads and abdomens were divided into pooled tissue samples for RNA extraction, cDNA synthesis, and qPCR analysis. Relative fold changes in gene expression are reported with respect to 0-24h PE well-fed form *pipiens* (Fig. 4–6). Statistical significance of fold change differences between forms for each time point were always examined by two-way ANOVA corrected for multiple comparisons. This also allowed us to assess significance of fold change differences between nutritional regimens within a bioform. With two discussed exceptions, we found no significant differences associated with diet treatment.

**Figure 4:**
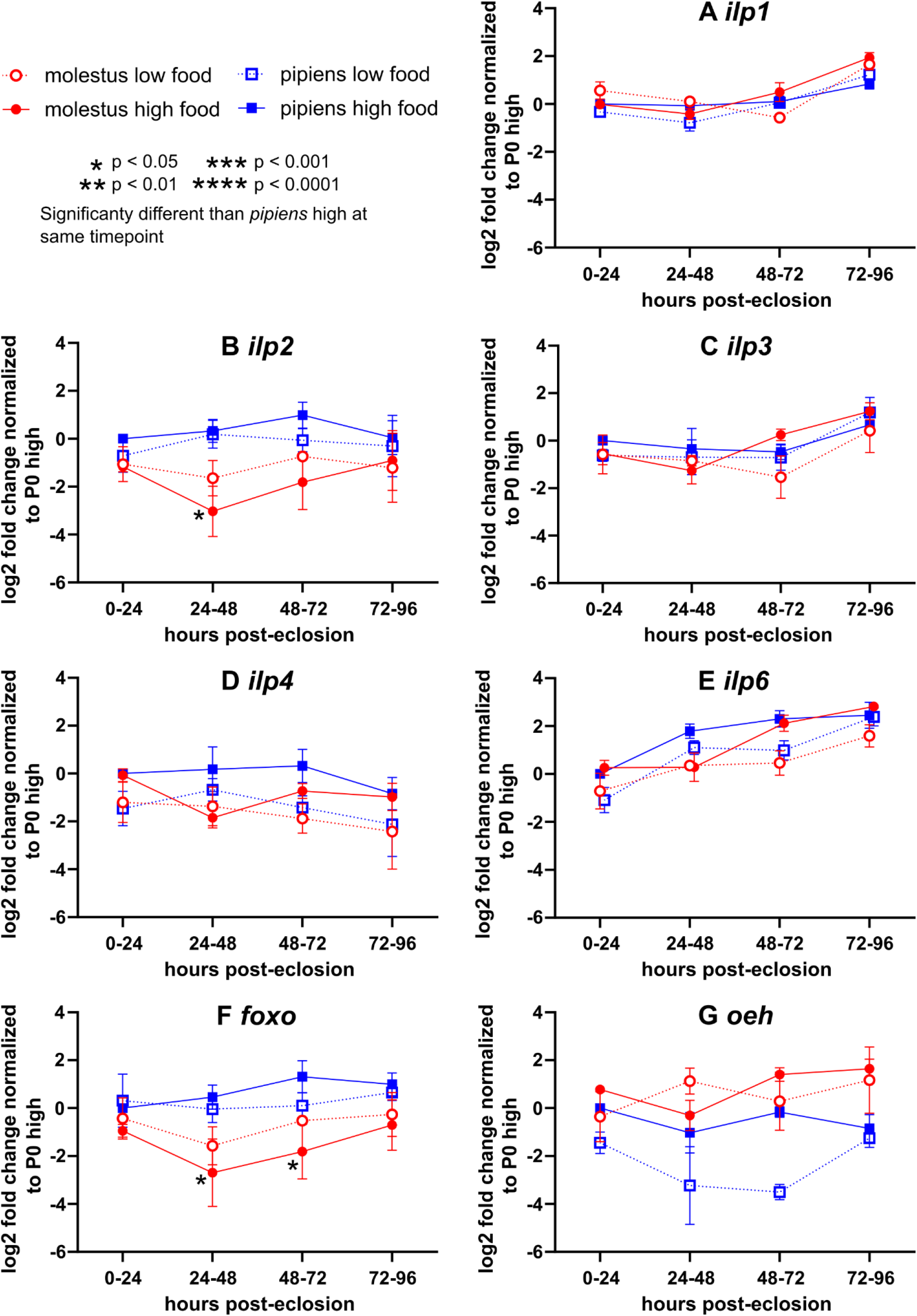
Downregulation of *ilp2* and *foxo* in head tissues are correlated with autogeny phenotype. qPCR was used to determine whether expression of *ilp1* (A), *ilp2* (B), *ilp3* (C), *ilp4* (D), *ilp6* (E), *foxo* (F), and *oeh* (G), differ between bioforms and feeding conditions over the first four days of adult development, when autogenous egg production occurs in *molestus*. Only *ilp2* (B) and *foxo* (F) are significantly differentially expressed, as determined by a two-way ANOVA of log2 fold change values. p values are indicated by *s as presented in the key. Fold change gene expression determined by ΔΔCt method, normalizing first to housekeeping gene *EF1a*, then to *pipiens* high food 0-24 hr reference sample. Data are shown on a log2 scale such that “0” means no change vs. the reference sample. “High” and “low” indicate diet types used in larval rearing. Error bars indicate SEM. Each data point is the mean of 3-4 biological replicates.

**Figure 5:**
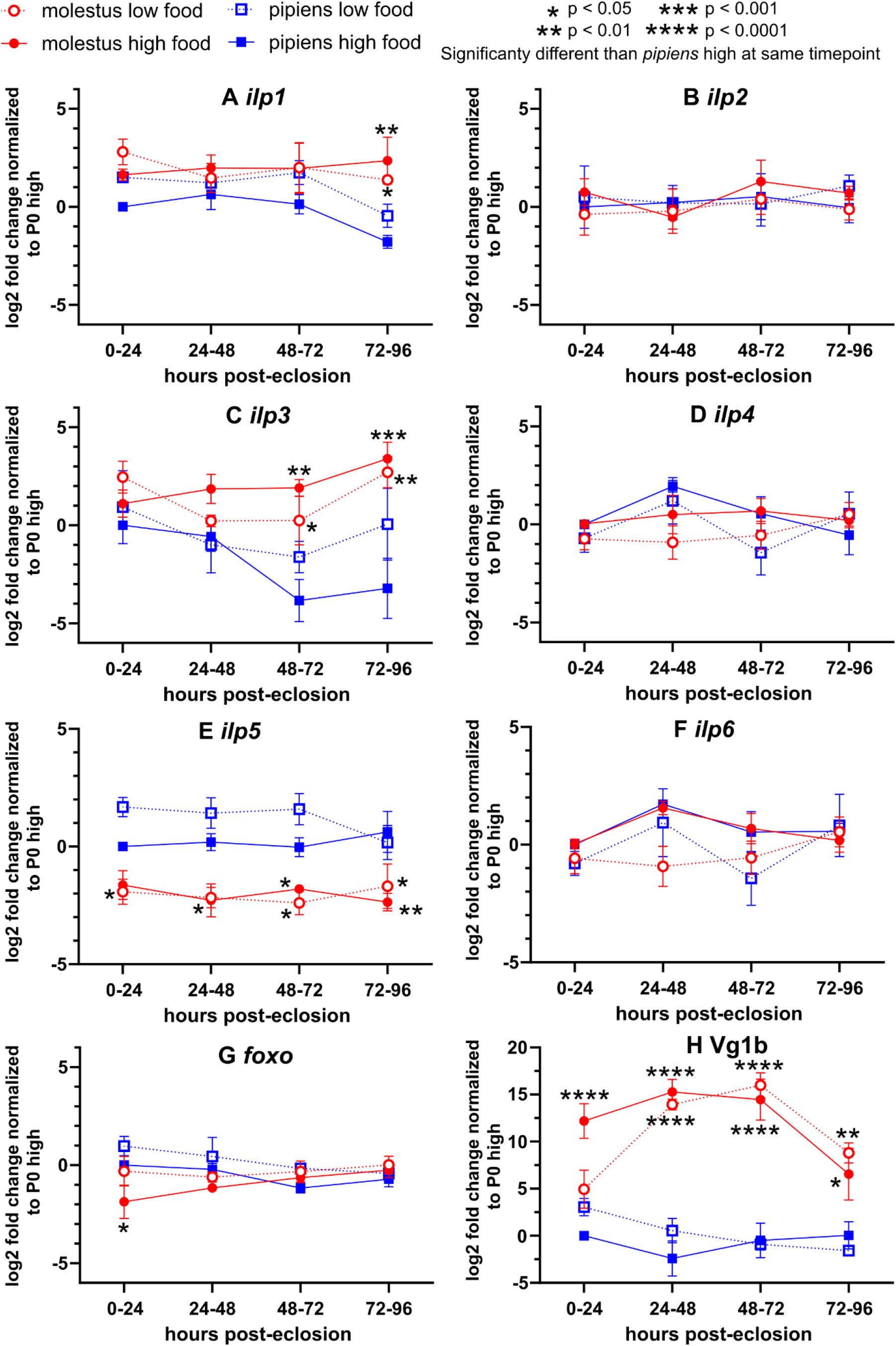
Multiple gene expression differences in the abdomen correlate with autogeny phenotype when biotypes and feeding conditions are compared. qPCR was used to determine whether expression of *ilp1* (A), *ilp2* (B), *ilp3* (C), *ilp4* (D), *ilp5* (E), *ilp6* (F), *foxo* (G), and *Vg1b* (H) differ between bioforms and feeding conditions over the first four days of adult development, when autogenous egg production occurs in *molestus. ilp1* is upregulated in *molestus* at 72-96 hrs post-eclosion (A). *ilp2* expression is stable between bioforms, feeding conditions and time points (B). *ilp3* is upregulated in molestus from 48-96 hrs post-eclosion (C). *ilp4* expression is stable between bioforms, feeding conditions and time points (D). *ilp5* is significantly downregulated at all timepoints in *molestus*, irrespective of feeding condition (E). *ilp6* expression is stable between bioforms, feeding conditions and time points (F). *foxo* is initially downregulated in well-fed *molestus* (G). *Vg1b* is enormously upregulated in molestus, but with a slight delay in onset in low-food conditions (H). p values are indicated by *s as presented in the key. Fold change gene expression determined by ΔΔCt method, normalizing first to housekeeping gene *EF1a*, then to *pipiens* high food 0-24 hr reference sample. Data are shown on a log2 scale such that “0” means no change vs. the reference sample. “High” and “low” indicate diet types used in larval rearing. Error bars indicate SEM. Each data point is the mean of 3-4 biological replicates.

**Figure 6:**
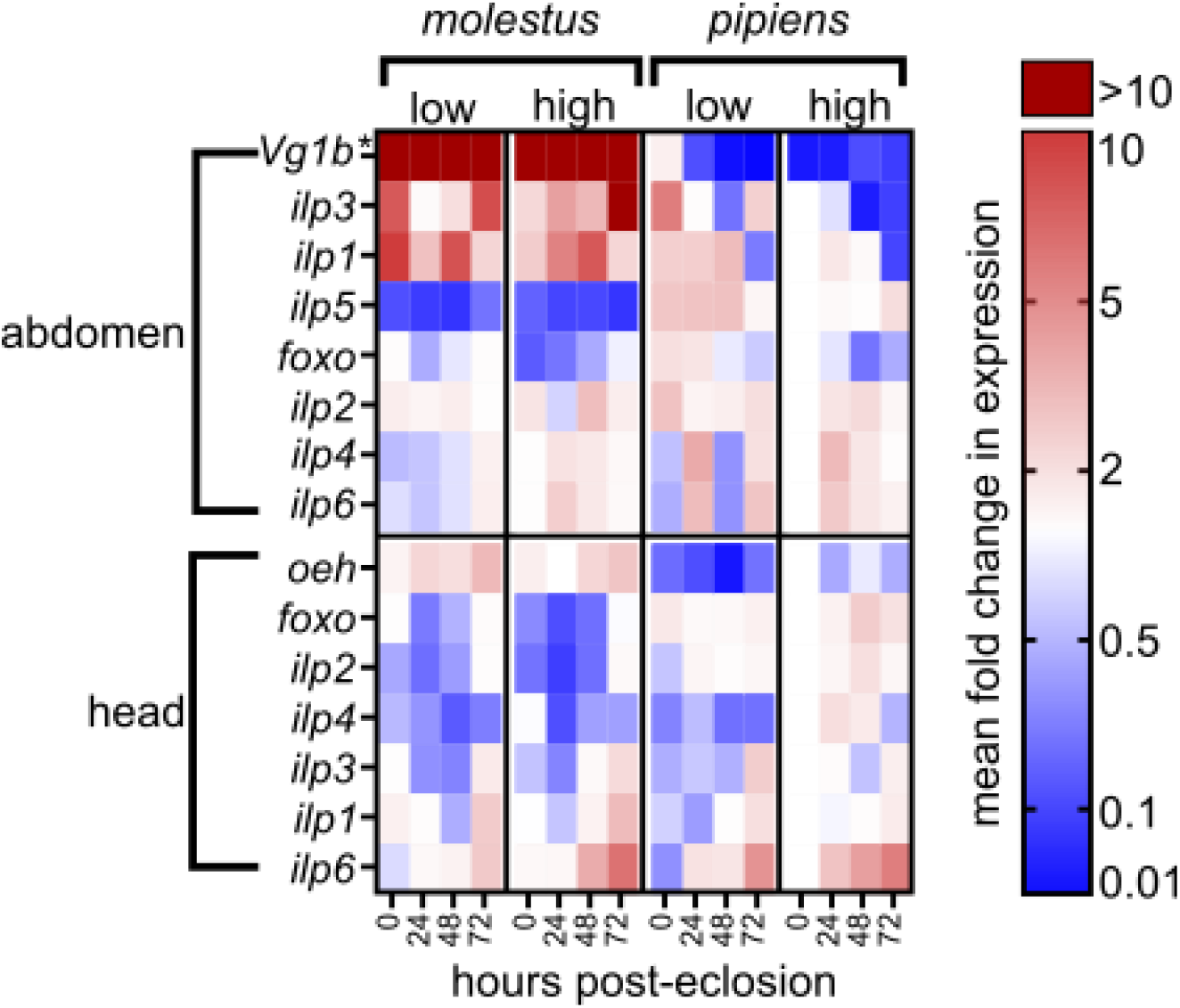
Heat Map Summary of Fold Change Gene Expression. Data from Figures 4 and 5 shown as fold changes in expression (as opposed to log2 scale in original figures) and are grouped to show genes that are up and downregulated during autogenic egg production (listed first in each grouping) vs. those that remain stable across forms (listed last in each grouping). *\*Vg1b* expression is shown as 1/100th of actual fold change values to allow for visualization on the same scale as other measured genes.

Previous work suggested that insulin signaling is an attractive candidate pathway for understanding the regulation of autogeny. For example, *ilp1* and *ilp5* are downregulated in diapausing *Cx. pipiens* females, which shuts down their ovarian maturation (Sim and Denlinger, 2009). RNAi knock-down of *ilp1* also results in cessation of ovarian development in females not programmed for diapause (Sim and Denlinger, 2009). Furthermore, ILP3 promotes ovarian maturation in both the autogenous *G. atropalpus* (Gulia-Nuss et al., 2012), as well as anautogenous *Ae. aegypti* following a blood meal (Brown et al., 2008). Brain medial neurosecretory cells are the dominant source of ILP expression not only in mosquitoes, but in insects in general (reviewed by Okamoto and Yamanaka, 2015). While ILP activity is controlled primarily by the release of these neuropeptides into the hemolymph, we predicted that genetically encoded regulatory changes between *Culex* bioforms could affect transcription levels, and subsequently, ILP activity. Therefore, we expected to see upregulation of *ilps*, especially *ilp3*, in the heads of *molestus* females, with the strongest upregulation occurring in those that were well-fed.

Females of both bioforms expressed *ilp1, ilp2, ilp3, ilp4* and *ilp6* in the head tissues, regardless of larval feeding conditions (Fig. 4A-E, Fig. 6). *ilp5* was the only insulin-like peptide that was undetectable in the head (data not shown). Expression of *ilp1* and *ilp6* increased over time but did so to the same extent in all groups, showing no correlation with autogenous egg production (Fig. 4A and E). Expression levels of *ilp1, ilp3, ilp4*, and *ilp6* did not differ between bioforms or feeding conditions in head tissues (Fig. 4A-E), even though ILP3 release from the brain is associated with mosquito reproduction in other systems (Brown et al., 2008). One exception was that *ilp1* expression was significantly different between well-fed and poorly-fed *molestus* at 48-72h PE (log2 = −0.57 vs. 0.5, p = 0.018)(Fig. 4A). This likely reflects a slight delay initiating upregulation of *ilp1* in poorly-fed *molestus*, which is ultimately upregulated in both diet treatments at 72-96h PE (log2 = 1.67 vs. 1.94). Interestingly, *ilp2* experienced a ten-fold downregulation in well-fed *molestus* at 24-48h PE vs. well-fed *pipiens* at the same time point (log2 = −3.03 vs. log2 = 0.33, p = 0.014), and more modest downregulation in poorly-fed *molestus* (log2 = −1.64)(Fig. 4B). The downstream effector of ILP signaling, *foxo*, was also detected in all head samples (Fig. 4F). Well-fed *molestus* downregulated *foxo* by nine-fold at 24-48h PE with respect to equivalently staged *pipiens* (log2 = −2.7 vs. 0.47, p = 0.021) and at 48-72h PE (log2 = −1.81 vs. 1.31, p = 0.023). Weaker downregulation of *foxo* was observed in poorly-fed *molestus* at those same time points (log2 = −1.57 and −0.52).

We also screened abdominal tissues as a potential alternative source of *ilp* expression. While not as well-known, there is a precedent for *ilp* expression originating from ovaries, fat body, and other abdominal tissues in many diverse insects (Okada et al., 2019; Okamoto et al., 2009a, 2009b). Insects can express ILPs directly from their fat bodies to signal nutritional conditions in a post-feeding stage. For example, in the beetle *Gnatocerus cornutus*, instead of the fat body signaling to the brain to synthesize and release ILPs in response to nutritional input, the fat body directly produces ILP to stimulate post-feeding growth (Okada et al., 2019). In mosquitoes, ILP expression of abdominal origin has also been characterized. For example, expression of ILP4 and ILP7 are detectable in ovary tissue in *Aedes* (Riehle et al., 2006). Furthermore, in female *Anopheles gambiae*, an antibody against ILP1/3/4 detected expression in abdominal ganglia that run along the body wall (Marquez et al., 2011). This same study identified ILP1/3/4 immunoreactivity in neuronal axons along the midgut and observed changes in the expression of several ILPs in the abdomen and midgut in response to sugar deprivation and exposure to human insulin via a bloodmeal. We reasoned that *ilp*s of similar abdominal or neural origin may be expressed in *molestus* prior to bloodfeeding and quantified gene expression in whole abdomens (Fig. 5). *ilp1* was significantly upregulated at 72-96h PE in both well and poorly-nourished *molestus* (log2 = 1.5 and 1.37 respectively) with respect to well-fed *pipiens* of the same age (log2 = −1.78) (p = 0.0014 vs. p = 0.02) (Fig. 5A). *ilp3* expression increased 53-fold in well-fed *molestus* compared to well-fed *pipiens* at 48-72h PE (log2 = 1.91 vs. −3.83, p = 0.003). The *ilp3* expression level differences between well-fed *molestus* and *pipiens* gradually rose to 97-fold by 72-96h PE (log2 = 3.39 vs. −3.21, p = 0.0002) (Fig. 5C). Poorly-nourished *molestus* significantly upregulated *ilp3* starting at 48-72h PE (16-fold, log2 = 0.24, p = 0.048), but expression levels did not reach those of well-fed *molestus* until 72h PE (60-fold, log2 = 2.71, p = 0.002). Oddly, the expression of *ilp3* in *molestus* was not significantly different when compared to poorly nourished *pipiens*. No differences in *ilp2, ilp4* or *ilp6* expression were observed between bioforms or feeding conditions (Fig. 5B-F). Finally, as was the case for head tissues, *foxo* expression was also briefly downregulated in the abdomens of well-fed *molestus*. We observed over three-fold lower expression in well-fed *molestus* vs. well-fed *pipiens* at 0-24h PE (log2 = −1.86 vs. 0, p = 0.040) (Fig. 5G).

Interestingly, *ilp5* was only expressed in abdominal tissues, and primarily by the *pipiens* bioform (Fig. 5E). Predominantly abdominal expression is conserved in *Aedes* and *Anopheles* mosquitoes (Krieger et al., 2004; Riehle et al., 2006), so it was unsurprising that *ilp5* was not detectable in head tissues. At any given time, expression was three to five times lower in *molestus* than well-fed *pipiens* of the same stage (log2 on average was approximately −2 vs 0, p < 0.05 in each case). While expression was equally low in *molestus* irrespective of nutritional regimen, in *pipiens*, poor nutrition was associated with modest, but not significant, upregulation of *ilp5*. These form and dietary differences in *ilp5* expression are especially intriguing. Previous work found that the expression of *ilp5* was associated with a non-diapause state in *Culex pipiens* (Sim and Denlinger, 2009). Indeed, one of the primary differences between the *pipiens* and *molestus* bioforms is the ability to undergo diapause (Denlinger and Armbruster, 2014; Vinogradova, 2000), although it is notable that knock-down of ILP5 had no effect on diapause phenotypes in previous work (Sim and Denlinger, 2009). Currently, the functional role of *ilp5* in *pipiens* is not well understood, but clues from *Aedes* suggest it is critical to acquisition of teneral reserves. For example, CRISPR-Cas9 ablation of *ilp5* in *Aedes* resulted in bigger mosquitoes with larger stores of lipids accumulated during larval stages (Ling and Raikhel, 2018). Because increases in female size and teneral reserves are critical for autogeny (Chambers and Klowden, 1994; Gulia-Nuss et al., 2015; Telang et al., 2006), the significant reduction in *ilp5* expression that we observed in the *molestus* bioform is potentially important for explaining a gain of autogenic ability.

To summarize our major findings for insulin signaling pathway genes, we observed brief downregulation of *ilp2* and *foxo* in the heads of females beginning at 24h PE. Upregulation of *ilp1* and *ilp3* occurred in the abdomens of *molestus* females after 48h PE, the latter of which experienced more dramatic upregulation. Abdominal expression of *foxo* was mildly downregulated in well-fed *molestus* females. Interestingly, *ilp5*, whose function is currently not well understood in *Cx. pipiens*, was only detected in the abdominal tissue, and strongly downregulated in *molestus* throughout the course of our experiment.

We then determined the time course of *Vg1b* expression to determine whether insulin-signaling pathway genes could be regulators of its expression. In anautogenous systems, *Vg* expression is repressed until a bloodmeal is acquired (Hansen et al., 2014). FOXO is the downstream effector of insulin signaling. It is a forkhead box transcription factor that represses the expression of *Vg* in many insects until the insulin signaling pathway is activated (Sheng et al., 2011). Insulin signaling triggers phosphorylation of FOXO and deportation from the nucleus to the cytoplasm, relieving the repression of *Vg* genes. In *molestus, Vg1b* was strongly upregulated (p < 0.0001) in abdominal tissues immediately upon eclosion in well-nourished samples and by 24-48h PE in poorly-nourished samples (Fig. 5H). Well-fed and poorly-fed *molestus* were significantly different at 0-24h PE (p = 0.011), owing to the delayed onset of upregulation that occurs in the poorly-fed sample. This agreed well with previous work in *Cx. tarsalis* which demonstrated robust upregulation of this gene within a day of emergence in autogenous females (Provost-Javier et al., 2010). Both well- and poorly-nourished *molestus* samples ultimately attained changes in expression greater than 65,000 times that of newly eclosed *pipiens*. It is notable that *Vg1b* expression peaked at 24-48h PE (log2 = 15.29) in well-fed *molestus* but was delayed until 48-72h PE (log2 = 16) in poorly-fed *molestus*. No upregulation occurs in *pipiens* in any feeding condition or at any time point (Fig. 5H). Interestingly, *Vg1b* is detected before adulthood in *Cx. tarsalis*, and in *Ae. albopictus, Vg* has an additional role in repressing blood-seeking behavior in young, sugar-fed females (Dittmer et al., 2019; Provost-Javier et al., 2010). We can infer that peak *Vg* transcription must occur before 48h PE because ovary maturation is visible by this point (Fig. 2). It may also be that very early upregulation of *Vg1b* serves to block blood-seeking behavior prior to or in addition to egg-producing functions.

We next examined expression patterns of insulin-signaling pathway genes in light of *Vg1b’s* temporal dynamics. An early-acting difference of potential importance for rapid *Vg1b* activation was the downregulation of *foxo* in 0-24h PE well-fed *molestus* abdomens. Downregulation of this conserved repressor of vitellogenesis genes could make it easier to activate *Vg1b* expression without bloodmeal-induced activation, even though it is typically regulated at the post-transcriptional level. However, the difference was only seen in well-fed *molestus* while autogeny occurred in both feeding regimens. Downregulation of *ilp5* also occurred early in *molestus*, but was observed in both diets tested, making it a better candidate regulator of *Vg1b*. While significantly upregulated in the abdomens of *molestus* females, *ilp1* and *ilp3* are not likely to be involved in upregulation of *Vg1b*. We did not observe form-specific expression differences at these genes until after autogenic egg production was in progress (Fig. 5A, 5C, Fig. 6). *ilp3* was not significantly upregulated until 48h PE in well-fed *molestus* and did not achieve a similar fold increase until 72h PE in poorly-fed *molestus*. By this point, *Vg1b* expression had passed its peak in both feeding conditions. Likewise, upregulation of *ilp1* specific to *molestus* did not occur until 72h PE. It may still reinforce the process once it is in progress since this *ilp* gene has already been shown to regulate ovarian development in *Culex* mosquitoes. Knock-down of expression of ILP1 has been demonstrated to halt ovarian development in non-diapausing *pipiens* and lower endogenous expression levels are observed in diapausing females (Sim and Denlinger, 2009). It is possible that changes in *ilp1* and *ilp3* expression have some important function in autogeny unrelated to *Vg1b* activation that will be determined by additional functional studies.

In *Aedes*, OEH, a neuroparsin type neuropeptide, activates the insulin signaling pathway in response to a bloodmeal, but bypasses the insulin receptor to do so (Dhara et al., 2013). More recent work demonstrated ecdysone signaling influences *ilp* expression in response to a bloodmeal, suggesting that it could also be important during autogeny (Ling and Raikhel, 2021). Based on previous predictions, we expected that OEH stimulation of egg production would be a conserved feature of mosquito reproduction (Vogel et al., 2015). We measured expression of *OEH* in the heads of females from both forms for 0-96h PE, with the expectation that it would be uniquely upregulated in *molestus* females. We did not detect expression in abdomens during pilot experiments. Although it was implicated in autogenous reproduction in other mosquito species (Brown et al., 2008; Dhara et al., 2013; Gulia-Nuss et al., 2015, 2012), *OEH* underwent very modest upregulation in the heads of *molestus* females throughout our time course relative to equivalently-staged *pipiens* (Fig. 4G, Fig. 6). OEH upregulation in the heads of *molestus* females preparing for autogenous egg production occurred after *Vg1b* upregulation and was not statistically significant compared to expression in equivalent-staged *pipiens* females (Fig. 4G, Fig. 6). In *G. atropalpus*, ecdysteroid is secreted by the ovaries immediately upon eclosion and peaks during the first day of emergence under high-food conditions, and experience a ~12 hour delay of peak secretion under low-food conditions (Telang et al., 2006). This is controlled by OEH from the brain released within 6 hours of eclosion and produced during pupation and the first few hours post-eclosion (Fuchs et al., 1980; Gulia-Nuss et al., 2012). It is possible that our wide 0-24 hour sampling window prevented detection of this early peak in OEH expression, or that OEH peak expression occurs during pupation for *molestus*. It may also be that OEH activity is predominantly regulated by release of peptide, and that transcriptional regulation is not a good indicator of its potential role in *molestus* autogeny.

Here, we characterized the effects of nutrition on time to pupation, autogenous egg production, and expression of genes connected to sensing of nutritional state in *Cx. pipiens*. We confirmed that genetic factors interacted with nutritional status to regulate autogenous egg production in *molestus*, while even well-fed *pipiens* never produced eggs autogenously. In agreement with this trait expression data, we observed strong and significant bioform-specific gene expression differences, which were only mildly influenced by nutrition. The comparison between autogenous *molestus* and anautogenous *pipiens* revealed changes in gene expression patterns in the abdomen (*ilp1, ilp3, ilp5*), head (*ilp2*), or both (*foxo*) (Fig. 6). Some of these coincided with dramatic upregulation of *Vg1b* (ex. *foxo*, *ilp5*), while others occurred afterward (ex. *ilp1, ilp3*). Gene expression differences in well- vs. poorly-nourished *molestus* could be characterized as dampened or delayed rather than statistically significant changes in expression, in agreement with incomplete loss of autogeny in our lowest diet treatment. Future work on the functional consequences of these gene expression differences, especially that of *ilp5*, will enhance our understanding of the evolution of autogeny and loss of diapause as bioform-specific traits.

## Data Availability

Scripts and data used to examine expression of autogeny under different dietary conditions can be found at https://github.com/mcadamme/Culex_Nutrition_Exp. Data from gene expression analysis can be found at: link to be added following acceptance.

## Experimental Procedures

### Gene identification and orthology

We identified six *Culex ilp* genes by performing a HMMR analysis in Vectorbase (https://legacy.vectorbase.org/hmmer) using an alignment of all eight *Drosophila* and *Aedes ilps* against the CpipJ2.4 geneset (Arensburger et al., 2010; Giraldo-Calderón et al., 2015). We next constructed phylogenetic trees to establish gene orthology. Putative *Culex* ILPs plus previously characterized insect ILP sequences were aligned in MUSCLE (https://www.ebi.ac.uk/Tools/msa/muscle/)(Edgar, 2004). Tree topology was determined by TOPALI v2.5 using both a MrBayes algorithm (Model: JTT + G, Runs: 2 Generations: 500000, Sample Freq.: 10, Burnin: 25%) and maximum PhyML algorithm (Model: JTT + G, 100 bootstrap runs)(Anisimova and Gascuel, 2006; Milne et al., 2004; Ronquist and Huelsenbeck, 2003). Synteny of *ilp* genes was examined using Genomicus Metazoa (web-code version: 2014-07-06, database version: 30.01, https://www.genomicus.biologie.ens.fr/genomicus-metazoa-30.01/cgi-bin/search.pl)(Louis et al., 2013).

### Mosquitoes

A below-ground population (form *molestus*) of *Cx. pipiens* was obtained from Calumet, IL, and has been in colony since 2009 (Mutebi and Savage, 2009) and maintained following Fritz et al. (Fritz et al., 2015, 2014). An above-ground population (form *pipiens*) of *Cx. pipiens* was obtained from Evanston, IL, and has been in colony since 2016 (Noreuil and Fritz 2021). Our laboratory-reared *molestus* had been cultured for 8 years without requiring a blood meal, but our *pipiens* never produced egg rafts prior to blood feeding during its 2 years in culture. All adults and larvae were maintained in an environmental chamber with a L:D photo period of 16h:8h, temperature of 25±1°C, and humidity of 50±10%. Adults for each form were kept in separate 60 × 60 × 60 cm white BugDorm-2 insect rearing cages (Megaview Science Education Services Co., Taichung, Taiwan) and were provided with a 10% sucrose solution at all times.

### Imaging of ovarian development

Pupae were picked from respective larval rearing pans, placed in individual tubes until eclosion, then moved to cages containing a 10% sucrose resource where females were allowed to age with or without males for one to four days. Ovaries were dissected under a stereomicroscope (Olympus Corporation, model SZ61, Center Valley, PA, USA) at 3.0-4.5 X magnification on a petri dish filled with 70% ethanol. Images were made using the Olympus cellSens Entry microscope-mounted imaging system.

### Impacts of genotype and nutrition on survivorship, larval development and ovarian maturation

To produce sufficient mosquitoes for experimentation, each colony was blood fed at 12-14 days of age using an artificial membrane feeder. Goose blood (Na-heparinated, obtained from Lampire Biological) sweetened with 50% sucrose solution (8:1 ratio) was provided through a pork sausage casing for a minimum of 1 hour. Approximately 72 hours after blood feeding females were provided with a cup of dechlorinated tap water and allowed to lay eggs for 48 hours. For each colony, the resulting egg rafts were placed in 27.0 × 19.4 × 9.5 cm plastic pans with 800ml of dechlorinated tap water and were supplied with 0.16g of liver powder (BD DifcoTM Desiccated, Powdered Beef Liver) and 0.08g of powdered inactive dry yeast (Genesee Scientific) suspended in 8ml of RO water. After 48 hours, 60 individuals from each colony were placed in one of four diet treatments using a 3mL plastic transfer pipette. Those treatments were as follows: extra low (0.25mg desiccated liver powder (LP), 0.14mg dry yeast (DY), 4ml RO water per larva), low (0.53mg LP, 0.27mg DY, 4ml RO water per larva), medium (0.80mg LP, 0.40mg DY, 4ml RO water per larva), and high (1.07mg LP, 0.53mg DY, 4ml RO water per larva). To avoid competition between individuals, larvae were placed in a single well of a 12-well culture plate. This resulted in 5 plates (12 × 5 = 60 individuals) per treatment, per form. Plates were held in an environmental chamber with the same conditions as described above. Individuals were fed every other day. Diet fed on days subsequent to the initial loading of the plates contained the same mass of food as described above but was suspended in 100μl of RO water instead of 4ml. Water was changed completely, and the diet was re-loaded as initially described at day 8 or 10 of the experiment to avoid build-up of waste within each well. The number of pupae present was recorded daily, and mortality was recorded every other day during feeding. Pupae were removed from the well and placed in a small plastic cup within a 30 × 30 × 30 white Bugdorm-2 cage. Adults were provided with a 10% sucrose solution and held under the previously described environmental conditions. Wing and ovarian dissections were performed on adult females, beginning at 4 days PE. A single wing was removed from each of 3 females per diet treatment, per form and photographed using an Olympus cellSens Entry microscope-mounted imaging system. Wing length measurements were made from the allular notch to the tip of the wing using ImageJ software (Schneider et al., 2012). Ovaries were also removed from at least 20 *molestus* females and 5 *pipiens* females per diet treatment, and the numbers of follicles advanced beyond Christopher’s stage IIb present were recorded (O’Meara and Krasnick, 1970). This experiment was repeated four times.

In a separate experiment, we measured the impacts of diet, form, and sex on larval development time. For each bioform and diet treatment (high or extra low), 60 larvae were placed into wells of the five 12-well cell culture plates and allowed to develop as described above. Upon pupation, individuals were transferred to a 2ml clear plastic lysis tube plugged with cotton wool. We monitored emergence every 48h and scored both larval development time and sex of the emerged adult. This experiment was repeated three times.

### Statistical analysis of mortality, pupation, and ovarian maturation

Bayesian generalized linear models (GLMs) were always fit using the package BRMS (v. 2.12.0 (Bürkner, 2017)) in R (R Core Team, 2016). Posterior probability distributions of model parameters were estimated using Markov Chain Monte Carlo (MCMC). For each model, we ran 4 chains with 6,000 steps, a burn-in of 2,000 steps, and saved every other step resulting in a total of 8,000 samples drawn from the posterior probability distribution. To ensure a stable sampling distribution had been reached, we visually examined trace plots, estimated effective sample size, and calculated Gelman and Rubin’s convergence diagnostic (Gelman and Rubin, 1992). Details of the models and their underlying distributions can be found in Tables S1-6, S8-11, and S13-16.

### RNA isolation and qPCR

To compare gene expression levels between species, developmental time points, and nutritional states, we first extracted total RNA from pools of tissue derived from ten like individuals using TRIzol Reagent (Invitrogen, Carlsbad, CA) according to the manufacturer’s instructions. Larvae from each bioform were reared individually in 12-well cell culture plates and fed either high or extra-low diet treatments. Pupae were placed in individual 2mL lysis tubes, allowed to emerge, and adult females were collected every 24 hours. From these females, replicate pools of ten heads (bearing intact chemosensory appendages) and ten abdomens were separately dissected on a small petri dish filled with dry ice for each diet treatment. Pooled tissue was stored in sterile 1.5 mL microcentrifuge tubes containing TRIzol Reagent at −80°C until RNA isolation was performed. One minor modification to this is that we back-extracted the aqueous RNA-containing layer with chloroform an additional time to remove trace amounts of TRIzol components that interfered with downstream steps. Once the resultant RNA pellet was dissolved in 20 μL of nuclease-free water, the RNA was treated to remove residual genomic DNA contamination with the Turbo DNA-free kit (Invitrogen, Carlsbad, CA) according to the manufacturer’s protocol. RNA yield was assessed following genomic DNA removal using a Nanodrop Lite spectrophotometer (Thermo Fisher Scientific, Waltham, MA). 250ng of RNA was used to synthesize cDNA with the iScript kit (BioRad, Hercules, CA). cDNA was diluted five-fold in nuclease-free water for use as a qPCR template, and total RNA was diluted to an equivalent dilution factor (2ng/μL) for –RT controls. qPCR was performed on a LightCycler 480 real-time PCR cycler (Roche, Basel, CH) using Luna Universal qPCR master mix (NEB, Ipswich, MA). Primer sequences can be found in Table S17. Results were analyzed by the ΔΔCt method (Livak and Schmittgen, 2001). Reference gene (*EF1A*) was selected using the Normfinder algorithm (Andersen et al., 2004) from a panel of six candidate reference genes selected from relevant literature (Ling and Salvaterra, 2011; Sim and Denlinger, 2009; Van Hiel et al., 2009). Statistical significance of qPCR data determined by 2-way ANOVA of ΔΔCts with Tukey’s multiple comparisons test using Graphpad software.

## Acknowledgements

We thank the University of Maryland’s Department of Cell Biology and Molecular Genetics Imaging Core for the use of their Roche LightCycler qPCR machine and Amy Beaven for expert training on the instrument. Thea Bliss conducted the wing measurements. We thank Leslie Pick (LP; UMD Dept of Entomology) for thoughtful discussions during the initial stages of this project. Kevin Vogel reviewed and provided constructive feedback on an early draft of our manuscript. AMCJ received support from NIH R01GM113230 to LP. This work was funded by University of Maryland startup funds and NIH R01AI125622A to MLF.

## References

Andersen, C.L., Jensen, J.L., Ørntoft, T.F., 2004. Normalization of Real-Time Quantitative Reverse Transcription-PCR Data: A Model-Based Variance Estimation Approach to Identify Genes Suited for Normalization, Applied to Bladder and Colon Cancer Data Sets. Cancer Res. 64, 5245–5250. https://doi.org/10.1158/0008-5472.CAN-04-0496

Anisimova, M., Gascuel, O., 2006. Approximate likelihood-ratio test for branches: A fast, accurate, and powerful alternative. Syst. Biol. 55, 539–552. https://doi.org/10.1080/10635150600755453

Arensburger, P., Megy, K., Waterhouse, R.M., Abrudan, J., Amedeo, P., Antelo, B., Bartholomay, L., Bidwell, S., Caler, E., Camara, F., Campbell, C.L., Campbell, K.S., Casola, C., Castro, M.T., Chandramouliswaran, I., Chapman, S.B., Christley, S., Costas, J., Eisenstadt, E., Feschotte, C., Fraser-Liggett, C., Guigo, R., Haas, B., Hammond, M., Hansson, B.S., Hemingway, J., Hill, S.R., Howarth, C., Ignell, R., Kennedy, R.C., Kodira, C.D., Lobo, N.F., Mao, C., Mayhew, G., Michel, K., Mori, A., Liu, N., Naveira, H., Nene, V., Nguyen, N., Pearson, M.D., Pritham, E.J., Puiu, D., Qi, Y., Ranson, H., Ribeiro, J.M.C., Roberston, H.M., Severson, D.W., Shumway, M., Stanke, M., Strausberg, R.L., Sun, C., Sutton, G., Tu, Z.J., Tubio, J.M.C., Unger, M.F., Vanlandingham, D.L., Vilella, A.J., White, O., White, J.R., Wondji, C.S., Wortman, J., Zdobnov, E.M., Birren, B., Christensen, B.M., Collins, F.H., Cornel, A., Dimopoulos, G., Hannick, L.I., Higgs, S., Lanzaro, G.C., Lawson, D., Lee, N.H., Muskavitch, M.A.T., Raikhel, A.S., Atkinson, P.W., 2010. Sequencing of Culex quinquefasciatus establishes a platform for mosquito comparative genomics. Science 330, 86–88. https://doi.org/10.1126/science.1191864

Aslamkhan, M., Laven, H., 1970. Inheritance of autogeny in the Culex pipiens complex. Pak. J. Zool. 2, 121–147.

Attardo, G.M., Hansen, I.A., Raikhel, A.S., 2005. Nutritional regulation of vitellogenesis in mosquitoes: Implications for anautogeny. Insect Biochem. Mol. Biol., Genetic manipulation of insects 35, 661–675. https://doi.org/10.1016/j.ibmb.2005.02.013

Brogiolo, W., Stocker, H., Ikeya, T., Rintelen, F., Fernandez, R., Hafen, E., 2001. An evolutionarily conserved function of the Drosophila insulin receptor and insulin-like peptides in growth control. Curr. Biol. CB 11, 213–221. https://doi.org/10.1016/s0960-9822(01)00068-9

Brown, M.R., Clark, K.D., Gulia, M., Zhao, Z., Garczynski, S.F., Crim, J.W., Suderman, R.J., Strand, M.R., 2008. An insulin-like peptide regulates egg maturation and metabolism in the mosquito Aedes aegypti. Proc. Natl. Acad. Sci. U. S. A. 105, 5716–5721. https://doi.org/10.1073/pnas.0800478105

Bürkner, P.-C., 2017. brms: An R Package for Bayesian Multilevel Models Using Stan. J. Stat. Softw. 80, 1–28. https://doi.org/10.18637/jss.v080.i01

Casasa, S., Moczek, A.P., 2018. Insulin signalling’s role in mediating tissue-specific nutritional plasticity and robustness in the horn-polyphenic beetle Onthophagus taurus. Proc. R. Soc. B Biol. Sci. 285, 20181631. https://doi.org/10.1098/rspb.2018.1631

Chambers, G.M., Klowden, M.J., 1994. Nutritional Reserves of Autogenous and Anautogenous Selected Strains of Aedes albopictus (Diptera: Culicidae). J. Med. Entomol. 31, 554–560. https://doi.org/10.1093/jmedent/31.4.554

Chandra, V., Fetter-Pruneda, I., Oxley, P.R., Ritger, A.L., McKenzie, S.K., Libbrecht, R., Kronauer, D.J.C., 2018. Social regulation of insulin signaling and the evolution of eusociality in ants. Science 361, 398–402. https://doi.org/10.1126/science.aar5723

Clifton, M.E., Noriega, F.G., 2012. The fate of follicles after a blood meal is dependent on previtellogenic nutrition and juvenile hormone in Aedes aegypti. J. Insect Physiol. 58, 1007–1019. https://doi.org/10.1016/j.jinsphys.2012.05.005

Colombani, J., Andersen, D.S., Léopold, P., 2012. Secreted peptide Dilp8 coordinates Drosophila tissue growth with developmental timing. Science 336, 582–585. https://doi.org/10.1126/science.1216689

Denlinger, D.L., Armbruster, P.A., 2014. Mosquito diapause. Annu. Rev. Entomol. 59, 73–93. https://doi.org/10.1146/annurev-ento-011613-162023

Dhara, A., Eum, J.-H., Robertson, A., Gulia-Nuss, M., Vogel, K.J., Clark, K.D., Graf, R., Brown, M.R., Strand, M.R., 2013. Ovary ecdysteroidogenic hormone functions independently of the insulin receptor in the yellow fever mosquito, Aedes aegypti. Insect Biochem. Mol. Biol. 43, 1100–1108. https://doi.org/10.1016/j.ibmb.2013.09.004

Dittmer, J., Alafndi, A., Gabrieli, P., 2019. Fat body–specific vitellogenin expression regulates host-seeking behaviour in the mosquito Aedes albopictus. PLOS Biol. 17, e3000238. https://doi.org/10.1371/journal.pbio.3000238

Edgar, R.C., 2004. MUSCLE: multiple sequence alignment with high accuracy and high throughput. Nucleic Acids Res. 32, 1792–1797. https://doi.org/10.1093/nar/gkh340

Feinsod, F.M., Spielman, A., 1980. Nutrient-mediated juvenile hormone secretion in mosquitoes. J. Insect Physiol. 26, 113–117. https://doi.org/10.1016/0022-1910(80)90050-5

Fonseca, D.M., Keyghobadi, N., Malcolm, C.A., Mehmet, C., Schaffner, F., Mogi, M., Fleischer, R.C., Wilkerson, R.C., 2004. Emerging vectors in the Culex pipiens complex. Science 303, 1535–1538. https://doi.org/10.1126/science.1094247

Fritz, M.L., Walker, E.D., Miller, J.R., Severson, D.W., Dworkin, I., 2015. Divergent host preferences of above- and below-ground Culex pipiens mosquitoes and their hybrid offspring. Med. Vet. Entomol. 29, 115–123. https://doi.org/10.1111/mve.12096

Fritz, M.L., Walker, E.D., Yunker, A.J., Dworkin, I., 2014. Daily blood feeding rhythms of laboratory-reared North American Culex pipiens. J. Circadian Rhythms 12, 1. https://doi.org/10.1186/1740-3391-12-1

Fuchs, M.S., Sundland, B.R., Kang, S.-H., 1980. In vivo induction of ovarian development in Aëdes atropalpus by a head extract from Aëdes aegypti. Int. J. Invertebr. Reprod. 2, 121–129. https://doi.org/10.1080/01651269.1980.10553347

Garelli, A., Gontijo, A.M., Miguela, V., Caparros, E., Dominguez, M., 2012. Imaginal discs secrete insulin-like peptide 8 to mediate plasticity of growth and maturation. Science 336, 579–582. https://doi.org/10.1126/science.1216735

Gelman, A., Rubin, D.B., 1992. Inference from Iterative Simulation Using Multiple Sequences. Stat. Sci. 7, 457–472. https://doi.org/10.1214/ss/1177011136

Giraldo-Calderón, G.I., Emrich, S.J., MacCallum, R.M., Maslen, G., Dialynas, E., Topalis, P., Ho, N., Gesing, S., VectorBase Consortium, Madey, G., Collins, F.H., Lawson, D., 2015. VectorBase: an updated bioinformatics resource for invertebrate vectors and other organisms related with human diseases. Nucleic Acids Res. 43, D707–713. https://doi.org/10.1093/nar/gku1117

Grönke, S., Clarke, D.-F., Broughton, S., Andrews, T.D., Partridge, L., 2010. Molecular Evolution and Functional Characterization of Drosophila Insulin-Like Peptides. PLOS Genet. 6, e1000857. https://doi.org/10.1371/journal.pgen.1000857

Gulia-Nuss, M., Elliot, A., Brown, M.R., Strand, M.R., 2015. Multiple factors contribute to anautogenous reproduction by the mosquito Aedes aegypti. J. Insect Physiol. 82, 8–16. https://doi.org/10.1016/j.jinsphys.2015.08.001

Gulia-Nuss, M., Eum, J.-H., Strand, M.R., Brown, M.R., 2012. Ovary ecdysteroidogenic hormone activates egg maturation in the mosquito Georgecraigius atropalpus after adult eclosion or a blood meal. J. Exp. Biol. 215, 3758–3767. https://doi.org/10.1242/jeb.074617

Hagedorn, H.H., O’Connor, J.D., Fuchs, M.S., Sage, B., Schlaeger, D.A., Bohm, M.K., 1975. The ovary as a source of alpha-ecdysone in an adult mosquito. Proc. Natl. Acad. Sci. U. S. A. 72, 3255–3259. https://doi.org/10.1073/pnas.72.8.3255

Hagedorn, H.H., Shapiro, J.P., Hanaoka, K., 1979. Ovarian ecdysone secretion is controlled by a brain hormone in an adult mosquito. Nature 282, 92–94. https://doi.org/10.1038/282092a0

Hansen, I.A., Attardo, G.M., Park, J.-H., Peng, Q., Raikhel, A.S., 2004. Target of rapamycin-mediated amino acid signaling in mosquito anautogeny. Proc. Natl. Acad. Sci. U. S. A. 101, 10626–10631. https://doi.org/10.1073/pnas.0403460101

Hansen, I.A., Attardo, G.M., Rodriguez, S.D., Drake, L.L., 2014. Four-way regulation of mosquito yolk protein precursor genes by juvenile hormone-, ecdysone-, nutrient-, and insulin-like peptide signaling pathways. Front. Physiol. 5. https://doi.org/10.3389/fphys.2014.00103

Harbach, R.E., Harbach, R.E., Harrison, B.A., Gad, A.M., 1984. Culex (Culex) Molestus Forskål (Diptera: Culicidae): neotype designation, description, variation, and taxonomic status. Proc. Entomol. Soc. Wash. 86, 521–542.

Kassim, N.F.A., Webb, C.E., Russell, R.C., 2012. Is the expression of autogeny by Culex molestus Forskal (Diptera: Culicidae) influenced by larval nutrition or by adult mating, sugar feeding, or blood feeding? J. Vector Ecol. 37, 162–171. https://doi.org/10.1111/j.1948-7134.2012.00213.x

Kent, R.J., Harrington, L.C., Norris, D.E., 2007. Genetic Differences Between Culex pipiens f. molestus and Culex pipiens pipiens (Diptera: Culicidae) in New York. J. Med. Entomol. 44, 50–59.

Krieger, M.J.B., Jahan, N., Riehle, M.A., Cao, C., Brown, M.R., 2004. Molecular characterization of insulin-like peptide genes and their expression in the African malaria mosquito, Anopheles gambiae. Insect Mol. Biol. 13, 305–315. https://doi.org/10.1111/j.0962-1075.2004.00489.x

Krishnamurthy, B.S., Laven, H., 1961. A Note on Inheritance of Autogeny in Culex Mosquitos. Bull. World Health Organ. 24, 675–677.

Ling, D., Salvaterra, P.M., 2011. Robust RT-qPCR Data Normalization: Validation and Selection of Internal Reference Genes during Post-Experimental Data Analysis. PLoS ONE 6. https://doi.org/10.1371/journal.pone.0017762

Ling, L., Raikhel, A.S., 2021. Cross-talk of insulin-like peptides, juvenile hormone, and 20-hydroxyecdysone in regulation of metabolism in the mosquito Aedes aegypti. Proc. Natl. Acad. Sci. 118. https://doi.org/10.1073/pnas.2023470118

Ling, L., Raikhel, A.S., 2018. Serotonin signaling regulates insulin-like peptides for growth, reproduction, and metabolism in the disease vector Aedes aegypti. Proc. Natl. Acad. Sci. 115, E9822–E9831. https://doi.org/10.1073/pnas.1808243115

Livak, K.J., Schmittgen, T.D., 2001. Analysis of relative gene expression data using real-time quantitative PCR and the 2(-Delta Delta C(T)) Method. Methods San Diego Calif 25, 402–408. https://doi.org/10.1006/meth.2001.1262

Louis, A., Muffato, M., Roest Crollius, H., 2013. Genomicus: five genome browsers for comparative genomics in eukaryota. Nucleic Acids Res. 41, D700–705. https://doi.org/10.1093/nar/gks1156

Lounibos, L.P., Van Dover, C., O’Meara, G.F., 1982. Fecundity, autogeny, and the larval environment of the pitcher-plant mosquito, Wyeomyia smithii. Oecologia 55, 160–164. https://doi.org/10.1007/BF00384482

Marquez, A.G., Pietri, J.E., Smithers, H.M., Nuss, A., Antonova, Y., Drexler, A.L., Riehle, M.A., Brown, M.R., Luckhart, S., 2011. Insulin-like peptides in the mosquito Anopheles stephensi: Identification and expression in response to diet and infection with Plasmodium falciparum. Gen. Comp. Endocrinol. 173, 303–312. https://doi.org/10.1016/j.ygcen.2011.06.005

Milne, I., Wright, F., Rowe, G., Marshall, D.F., Husmeier, D., McGuire, G., 2004. TOPALi: software for automatic identification of recombinant sequences within DNA multiple alignments. Bioinforma. Oxf. Engl. 20, 1806–1807. https://doi.org/10.1093/bioinformatics/bth155

Mori, A., Romero-Severson, J., Black, W.C., Severson, D.W., 2008. Quantitative trait loci determining autogeny and body size in the Asian tiger mosquito (Aedes albopictus). Heredity 101, 75–82. https://doi.org/10.1038/hdy.2008.32

Mostowy, W., Foster, W., 2004. Antagonistic effects of energy status on meal size and egg-batch size of Aedes aegypti (Diptera: Culicidae). J. Vector Ecol. J. Soc. Vector Ecol. 29, 84–93.

Mutebi, J.-P., Savage, H.M., 2009. Discovery of Culex pipiens pipiens form molestus in Chicago. J. Am. Mosq. Control Assoc. 25, 500–503. https://doi.org/10.2987/09-5910.1

Nijhout, H.F., McKenna, K.Z., 2018. The distinct roles of insulin signaling in polyphenic development. Curr. Opin. Insect Sci. 25, 58–64. https://doi.org/10.1016/j.cois.2017.11.011

Noriega, F.G., 2004. Nutritional regulation of JH synthesis: a mechanism to control reproductive maturation in mosquitoes? Insect Biochem. Mol. Biol., Molecular and population biology of mosquitoes 34, 687–693. https://doi.org/10.1016/j.ibmb.2004.03.021

Okada, Y., Katsuki, M., Okamoto, N., Fujioka, H., Okada, K., 2019. A specific type of insulin-like peptide regulates the conditional growth of a beetle weapon. PLOS Biol. 17, e3000541. https://doi.org/10.1371/journal.pbio.3000541

Okamoto, N., Yamanaka, N., 2015. Nutrition-dependent control of insect development by insulin-like peptides. Curr. Opin. Insect Sci., Global change biology * Molecular physiology 11, 21–30. https://doi.org/10.1016/j.cois.2015.08.001

Okamoto, N., Yamanaka, N., Satake, H., Saegusa, H., Kataoka, H., Mizoguchi, A., 2009a. An ecdysteroid-inducible insulin-like growth factor-like peptide regulates adult development of the silkmoth Bombyx mori. FEBS J. 276, 1221–1232. https://doi.org/10.1111/j.1742-4658.2008.06859.x

Okamoto, N., Yamanaka, N., Yagi, Y., Nishida, Y., Kataoka, H., O’Connor, M.B., Mizoguchi, A., 2009b. A fat body-derived IGF-like peptide regulates postfeeding growth in Drosophila. Dev. Cell 17, 885–891. https://doi.org/10.1016/j.devcel.2009.10.008

O’Meara, G.F., 1985. Gonotrophic Interactions in Mosquitoes: Kicking the Blood-Feeding Habit. Fla. Entomol. 68, 122–133. https://doi.org/10.2307/3494335

O’meara, G.F., Jr, C., B, G., 1969. Monofactorial inheritance of autogeny in Aedes atropalpus. Mosq. News 29.

O’Meara, G.F., Krasnick, G.J., 1970. Dietary and Genetic Control of the Expression of Autogenous Reproduction in Aedes Atropalpus (Coq.) (Diptera: Culicidae)1. J. Med. Entomol. 7, 328–334. https://doi.org/10.1093/jmedent/7.3.328

Provost-Javier, K.N., Chen, S., Rasgon, J.L., 2010. Vitellogenin gene expression in autogenous Culex tarsalis. Insect Mol. Biol. 19, 423–429. https://doi.org/10.1111/j.1365-2583.2010.00999.x

R Core Team, 2016. R: A language and environment for statistical computing. R Foundation for Statistical Computing.

Riehle, M.A., Fan, Y., Cao, C., Brown, M.R., 2006. Molecular characterization of insulin-like peptides in the yellow fever mosquito, Aedes aegypti: expression, cellular localization, and phylogeny. Peptides 27, 2547–2560. https://doi.org/10.1016/j.peptides.2006.07.016

Riehle, M.A., Garczynski, S.F., Crim, J.W., Hill, C.A., Brown, M.R., 2002. Neuropeptides and Peptide Hormones in Anopheles gambiae. Science 298, 172–175. https://doi.org/10.1126/science.1076827

Ronquist, F., Huelsenbeck, J.P., 2003. MrBayes 3: Bayesian phylogenetic inference under mixed models. Bioinformatics 19, 1572–1574. https://doi.org/10.1093/bioinformatics/btg180

Roubaud, E., 1929. Autogenous Cycle of Winter Generations of Culex pipiens L. C. r. Acad. Sci. 188.

Roy, S., Smykal, V., Johnson, L., Saha, T.T., Zou, Z., Raikhel, A.S., 2016. Chapter Five - Regulation of Reproductive Processes in Female Mosquitoes, in: Raikhel, Alexander S. (Ed.), Advances in Insect Physiology, Progress in Mosquito Research. Academic Press, pp. 115–144. https://doi.org/10.1016/bs.aiip.2016.05.004

Roy, S.G., Hansen, I.A., Raikhel, A.S., 2007. Effect of insulin and 20-hydroxyecdysone in the fat body of the yellow fever mosquito, Aedes aegypti. Insect Biochem. Mol. Biol. 37, 1317–1326. https://doi.org/10.1016/j.ibmb.2007.08.004

Schneider, C. A., Rasband, W. S., Eliceiri, K. W., 2012. NIH Image to ImageJ: 25 years of image analysis. Nat. Methods, 9(7), 671–675.

Sharma, A., Nuss, A.B., Gulia-Nuss, M., 2019. Insulin-Like Peptide Signaling in Mosquitoes: The Road Behind and the Road Ahead. Front. Endocrinol. 10. https://doi.org/10.3389/fendo.2019.00166

Sheng, Z., Xu, J., Bai, H., Zhu, F., Palli, S.R., 2011. Juvenile Hormone Regulates Vitellogenin Gene Expression through Insulin-like Peptide Signaling Pathway in the Red Flour Beetle, Tribolium castaneum. J. Biol. Chem. 286, 41924–41936. https://doi.org/10.1074/jbc.M111.269845

Sim, C., Denlinger, D.L., 2009. A shut-down in expression of an insulin-like peptide, ILP-1, halts ovarian maturation during the overwintering diapause of the mosquito Culex pipiens. Insect Mol. Biol. 18, 325–332. https://doi.org/10.1111/j.1365-2583.2009.00872.x

Spielman, A., 2001. Structure and seasonality of nearctic Culex pipiens populations. Ann. N. Y. Acad. Sci. 951, 220–234. https://doi.org/10.1111/j.1749-6632.2001.tb02699.x

Spielman, A., 1957. The inheritance of autogeny in the Culex pipiens complex of mosquitoes.. Am. J. Epidemiol. 65, 404–425. https://doi.org/10.1093/oxfordjournals.aje.a119878

Spielman, A., Wong, J., 1973. Environmental Control of Ovarian Diapause in Culex pipiens. https://doi.org/10.1093/AESA/66.4.905

Telang, A., Li, Y., Noriega, F.G., Brown, M.R., 2006. Effects of larval nutrition on the endocrinology of mosquito egg development. J. Exp. Biol. 209, 645–655. https://doi.org/10.1242/jeb.02026

Telang, A., Rechel, J.A., Brandt, J.R., Donnell, D.M., 2013. Analysis of ovary-specific genes in relation to egg maturation and female nutritional condition in the mosquitoes Georgecraigius atropalpus and Aedes aegypti (Diptera: Culicidae). J. Insect Physiol. 59, 283–294. https://doi.org/10.1016/j.jinsphys.2012.11.006

Telang, A., Wells, M.A., 2004. The effect of larval and adult nutrition on successful autogenous egg production by a mosquito. J. Insect Physiol. 50, 677–685. https://doi.org/10.1016/j.jinsphys.2004.05.001

Trpis, M., 1978. Genetics of hematophagy and autogeny in the Aedes scutellaris complex (Diptera: Culicidae). J. Med. Entomol. 15, 73–80. https://doi.org/10.1093/jmedent/15.1.73

Tsuji, N., Okazawa, T., Yamamura, N., 1990. Autogenous and Anautogenous Mosquitoes: a Mathematical Analysis of Reproductive Strategies. J. Med. Entomol. 27, 446–453. https://doi.org/10.1093/jmedent/27.4.446

Van Hiel, M.B., Van Wielendaele, P., Temmerman, L., Van Soest, S., Vuerinckx, K., Huybrechts, R., Broeck, J.V., Simonet, G., 2009. Identification and validation of housekeeping genes in brains of the desert locust Schistocerca gregaria under different developmental conditions. BMC Mol. Biol. 10, 56. https://doi.org/10.1186/1471-2199-10-56

Vinogradova, A.B., 2000. Culex Pipiens Pipiens Mosquitoes: Taxonomy, Distribution, Ecology, Physiology, Genetics, Applied Importance and Control. Pensoft Publishers, Sofia.

Vogel, K.J., Brown, M.R., Strand, M.R., 2015. Ovary ecdysteroidogenic hormone requires a receptor tyrosine kinase to activate egg formation in the mosquito Aedes aegypti. Proc. Natl. Acad. Sci. 112, 5057–5062. https://doi.org/10.1073/pnas.1501814112

Wheeler, D.E., Frederik Nijhout, H., 1983. Soldier determination in Pheidole bicarinata: Effect of methoprene on caste and size within castes. J. Insect Physiol. 29, 847–854. https://doi.org/10.1016/0022-1910(83)90151-8

Yurchenko, A.A., Masri, R.A., Khrabrova, N.V., Sibataev, A.K., Fritz, M.L., Sharakhova, M.V., 2020. Genomic differentiation and intercontinental population structure of mosquito vectors Culex pipiens pipiens and Culex pipiens molestus. Sci. Rep. 10. https://doi.org/10.1038/s41598-020-63305-z

Zou, Z., Saha, T.T., Roy, S., Shin, S.W., Backman, T.W.H., Girke, T., White, K.P., Raikhel, A.S., 2013. Juvenile hormone and its receptor, methoprene-tolerant, control the dynamics of mosquito gene expression. Proc. Natl. Acad. Sci. https://doi.org/10.1073/pnas.1305293110

