## Supplemental Files for "Autogenous and anautogenous *Culex pipiens* bioforms exhibit insulin-like peptide signaling pathway gene expression differences that are not dependent upon larval nutrition"

**Supporting Information - Jarvela *et al.***

*Supplemental Tables*

Table S1: Estimates of beta coefficients from Bayesian generalized linear model with a binomial distribution quantifying the relationship between mortality, diet treatment, strain, diet treatment * strain and start date. Number Alive | Number of trials (alive + dead) ~ diet treatment + strain + diet treatment * strain + start date. Overlap of credible intervals with 0 indicates the beta coefficients (effect sizes) are not different from zero.

| **Model Term** | **Estimate of beta coefficient** | **Estimated error** | **Lower 95% credible interval** | **Upper 95% credible interval** |
|---|---|---|---|---|
| Intercept (high diet treatment, molestus, start date 1) | 2.06 | 0.26 | 1.57 | 2.57 |
| Medium diet | -0.01 | 0.33 | -0.67 | 0.64 |
| Low diet | 0.07 | 0.34 | -0.59 | 0.75 |
| Extra low diet | -0.51 | 0.31 | -1.12 | 0.08 |
| Start date 2 | 0.54 | 0.20 | 0.15 | 0.94 |
| Start date 3 | 0.53 | 0.21 | 0.13 | 0.93 |
| Start date 4 | 0.53 | 0.21 | 0.13 | 0.94 |
| Strain pipiens | -0.52 | 0.31 | -1.13 | 0.06 |
| Medium diet: Strain pipiens | 0.28 | 0.44 | -0.58 | 1.15 |
| Low diet: Strain pipiens | 0.00 | 0.44 | -0.87 | 0.87 |
| Extra low diet: Strain pipiens | 0.33 | 0.41 | -0.45 | 1.14 |

Table S2: Rhat and effective sample sizes for Bayesian generalized linear model with a binomial distribution exploring the relationship between mortality, diet treatment, strain, diet treatment * strain and start date. Number Alive | Number of trials (alive + dead) ~ diet treatment + strain + diet treatment * strain + start date

| **Model Term** | **Rhat** | **Bulk Effective Sample Size** | **Tail Effective Sample Size** |
|---|---|---|---|
| Intercept (high diet treatment, molestus, start date 1) | 1.00 | 5875 | 6655 |
| Medium diet | 1.00 | 5770 | 6182 |
| Low diet | 1.00 | 5778 | 6257 |
| Extra low diet | 1.00 | 6180 | 6522 |
| Start date 2 | 1.00 | 7690 | 7392 |
| Start date 3 | 1.00 | 7711 | 7439 |
| Start date 4 | 1.00 | 7738 | 7358 |
| Strain pipiens | 1.00 | 5504 | 6372 |
| Medium diet: Strain pipiens | 1.00 | 5623 | 6342 |
| Low diet: Strain pipiens | 1.00 | 5775 | 6379 |
| Extra low diet: Strain pipiens | 1.00 | 5847 | 6274 |

Table S3: Estimates of beta coefficients from Bayesian generalized linear model with a gaussian distribution quantifying the relationship between wing length, diet treatment, form, and start date. Wing length ~ diet treatment + strain + start date. Overlap of credible intervals with 0 indicates the beta coefficients (effect sizes) are not different from zero.

| **Model Term** | **Estimate of beta coefficient** | **Estimated error** | **Lower 95% credible interval** | **Upper 95% credible interval** |
|---|---|---|---|---|
| Intercept (high diet treatment, molestus, start date 1) | 3.51 | 0.04 | 3.43 | 3.58 |
| Medium diet | -0.18 | 0.04 | -0.26 | -0.11 |
| Low diet | -0.36 | 0.04 | -0.43 | -0.28 |
| Extra low diet | -0.55 | 0.04 | -0.62 | -0.47 |
| Start date 2 | -0.13 | 0.04 | -0.21 | -0.06 |
| Start date 3 | -0.11 | 0.04 | -0.18 | -0.03 |
| Start date 4 | -0.15 | 0.04 | -0.22 | -0.07 |
| Strain pipiens | 0.28 | 0.03 | 0.22 | 0.33 |

Table S4: Rhat and effective sample sizes for Bayesian generalized linear model with a gaussian distribution quantifying the relationship between wing length, diet treatment, form, and start date. Wing length ~ diet treatment + strain + start date.

| **Model Term** | **Rhat** | **Bulk Effective Sample Size** | **Tail Effective Sample Size** |
|---|---|---|---|
| Intercept (high diet treatment, molestus, start date 1) | 1.00 | 7471 | 6256 |
| Medium diet | 1.00 | 6707 | 7606 |
| Low diet | 1.00 | 6975 | 6728 |
| Extra low diet | 1.00 | 7512 | 7354 |
| Start date 2 | 1.00 | 7216 | 6789 |
| Start date 3 | 1.00 | 7197 | 7282 |
| Start date 4 | 1.00 | 7053 | 6977 |
| Strain pipiens | 1.00 | 7477 | 6837 |

Table S5: Estimates of beta coefficients from Bayesian generalized linear model with a Poisson distribution quantifying the relationship between the number of days until pupation, diet treatment, strain, diet treatment * strain and start date. Days to Pupation ~ diet treatment + strain + diet treatment * strain + start date. Overlap of credible intervals with 0 indicates the beta coefficients (effect sizes) are not different from zero. We transformed data whereby the first day pupae began to emerge is coded as day 0 (which corresponds to day 6 total experiment time).

| **Model Term** | **Estimate of beta coefficient** | **Estimated error** | **Lower 95% credible interval** | **Upper 95% credible interval** |
|---|---|---|---|---|
| Intercept (high diet treatment, molestus, start date 1) | 0.69 | 0.06 | 0.58 | 0.80 |
| Medium diet | 0.16 | 0.07 | 0.02 | 0.29 |
| Low diet | 0.49 | 0.06 | 0.37 | 0.62 |
| Extra low diet | 1.38 | 0.06 | 1.27 | 1.49 |
| Strain pipiens | -0.39 | 0.08 | -0.54 | -0.23 |
| Start date 2 | 0.10 | 0.04 | 0.03 | 0.18 |
| Start date 3 | -0.32 | 0.04 | -0.40 | -0.23 |
| Start date 4 | -0.34 | 0.04 | -0.42 | -0.26 |
| Medium diet: Strain pipiens | -0.01 | 0.11 | -0.22 | 0.21 |
| Low diet: Strain pipiens | -0.07 | 0.10 | -0.28 | 0.13 |
| Extra low diet: Strain pipiens | -0.14 | 0.09 | -0.32 | 0.04 |

Table S6: Rhat and effective sample sizes for Bayesian generalized linear model with a truncated Poisson distribution exploring the relationship between the number of days until pupation, diet treatment, strain, diet treatment * strain and start date. Days to Pupation ~ diet treatment + strain + diet treatment * strain + start date

| **Model Term** | **Rhat** | **Bulk Effective Sample Size** | **Tail Effective Sample Size** |
|---|---|---|---|
| Intercept (high diet treatment, molestus, start date 1) | 1.00 | 5323 | 6187 |
| Medium diet | 1.00 | 5505 | 6677 |
| Low diet | 1.00 | 5276 | 6859 |
| Extra low diet | 1.00 | 5231 | 6648 |
| Strain pipiens | 1.00 | 5022 | 6636 |
| Start date 2 | 1.00 | 7403 | 7164 |
| Start date 3 | 1.00 | 7618 | 7147 |
| Start date 4 | 1.00 | 7656 | 7540 |
| Medium diet: Strain pipiens | 1.00 | 5217 | 6048 |
| Low diet: Strain pipiens | 1.00 | 5346 | 6153 |
| Extra low diet: Strain pipiens | 1.00 | 4898 | 6298 |

Table S7: Summary of mean number of days until pupation, standard deviation, and total number of pupae per diet treatment, per replicate, per strain.

| **Diet Treatment** | **Start Date** | **Mean Molestus** | **Std. Dev. Molestus** | **Number Molestus** | **Mean Pipiens** | **Std. Dev. Pipiens** | **Number Pipiens** |
|---|---|---|---|---|---|---|---|
| Extra Low | 1 | 12.68 | 2.43 | 44 | 9.57 | 1.1 | 49 |
| | 2 | 14.02 | 2.15 | 54 | 11.65 | 1.73 | 51 |
| | 3 | 13.63 | 3.38 | 49 | 9.73 | 2.23 | 49 |
| | 4 | 11.83 | 1.76 | 53 | 9.74 | 1.35 | 53 |
| | **All** | **13.05** | **2.62** | **200** | **10.18** | **1.85** | **202** |
| Low | 1 | 9.42 | 1.23 | 52 | 8.12 | 0.64 | 46 |
| | 2 | 9.51 | 1.42 | 51 | 8.82 | 1.50 | 55 |
| | 3 | 8.46 | 1.65 | 56 | 6.7 | 0.77 | 53 |
| | 4 | 8.2 | 1.07 | 56 | 7.67 | 0.64 | 55 |
| | **All** | **8.87** | **1.47** | **215** | **7.82** | **1.24** | **209** |
| Medium | 1 | 9.12 | 1.04 | 52 | 7.86 | 0.7 | 50 |
| | 2 | 8.57 | 0.99 | 56 | 8 | 1.18 | 57 |
| | 3 | 7.53 | 0.68 | 59 | 6.38 | 0.56 | 55 |
| | 4 | 7.13 | 0.76 | 53 | 7.4 | 0.6 | 53 |
| | **All** | **8.07** | **1.17** | **220** | **7.4** | **1.03** | **215** |
| High | 1 | 8.66 | 0.62 | 53 | 7.65 | 0.60 | 46 |
| | 2 | 8.2 | 0.1 | 56 | 7.25 | 0.8 | 55 |
| | 3 | 7.54 | 0.69 | 56 | 6.63 | 0.76 | 54 |
| | 4 | 6.75 | 0.71 | 57 | 7.36 | 0.56 | 53 |
| | **All** | **7.77** | **1.05** | **222** | **7.21** | **0.78** | **208** |

Table S8: Estimates of beta coefficients from Bayesian generalized linear model with a Poisson distribution quantifying the relationship between larval development time, form, sex, diet treatment (extra low v. high), and start date. Days to Pupation ~ diet treatment + strain + sex + diet treatment * strain + diet treatment * sex + sex * strain + diet treatment * strain * sex + start date. Overlap of credible intervals with 0 indicates the beta coefficients (effect sizes) are not different from zero.

| **Model Term** | **Estimate of beta coefficient** | **Estimated error** | **Lower 95% credible interval** | **Upper 95% credible interval** |
|---|---|---|---|---|
| Intercept (high diet treatment, molestus, female, start date 1) | 0.62 | 0.04 | 0.53 | 0.70 |
| Extra low diet | 1.54 | 0.05 | 1.45 | 1.63 |
| Male | -0.46 | 0.07 | -0.60 | -0.34 |
| Strain pipiens | -0.53 | 0.07 | -0.68 | -0.39 |
| Start date 2 | -0.02 | 0.03 | -0.08 | 0.03 |
| Start date 3 | 0.05 | 0.02 | 0 | 0.10 |
| Extra low diet: Male | -0.12 | 0.07 | -0.26 | 0.02 |
| Extra low diet: Strain pipiens | 0.12 | 0.08 | -0.04 | 0.27 |
| Male: Strain pipiens | 0.26 | 0.11 | 0.05 | 0.47 |
| Extra low diet: Male: Strain Pipiens | -0.36 | 0.12 | -0.59 | -0.13 |

Table S9: Rhat and effective sample sizes for Bayesian generalized linear model with a a Poisson distribution quantifying the relationship between larval development time, form, sex, diet treatment (extra low v. high), and start date. Days to Pupation ~ diet treatment + strain + sex + diet treatment * strain + diet treatment * sex + sex * strain + diet treatment * strain * sex + start date.

| **Model Term** | **Rhat** | **Bulk Effective Sample Size** | **Tail Effective Sample Size** |
|---|---|---|---|
| Intercept (high diet treatment, molestus, female, start date 1) | 1.00 | 6928 | 7202 |
| Extra low diet | 1.00 | 6477 | 7205 |
| Male | 1.00 | 6170 | 6875 |
| Strain pipiens | 1.00 | 5760 | 6352 |
| Start date 2 | 1.00 | 7283 | 6684 |
| Start date 3 | 1.00 | 7846 | 7116 |
| Extra low diet: Male | 1.00 | 6113 | 6949 |
| Extra low diet: Strain pipiens | 1.00 | 5849 | 6142 |
| Male: Strain pipiens | 1.00 | 5103 | 6453 |
| Extra low diet: Male: Strain Pipiens | 1.00 | 5065 | 5970 |

Table S10: Estimates of beta coefficients from Bayesian generalized linear model with a zero-inflated Poisson distribution quantifying the relationship between the number of elongated follicles, diet treatment, and start date. Number of follicles ~ diet treatment + start date. Overlap of credible intervals with 0 indicates the beta coefficients (effect sizes) are not different from zero.

| **Model Term** | **Estimate of beta coefficient** | **Estimated error** | **Lower 95% credible interval** | **Upper 95% credible interval** |
|---|---|---|---|---|
| Intercept (high diet treatment, start date 1) | 3.96 | 0.03 | 3.91 | 4.02 |
| Medium diet | -0.18 | 0.03 | -0.23 | -0.12 |
| Low diet | -0.52 | 0.03 | -0.57 | -0.46 |
| Extra low diet | -1.00 | 0.04 | -1.08 | -0.92 |
| Start date 2 | -0.10 | 0.03 | -0.16 | -0.04 |
| Start date 3 | -0.06 | 0.03 | -0.12 | -0.00 |
| Start date 4 | -0.17 | 0.03 | -0.24 | -0.11 |

Table S11: Rhat and effective sample sizes for Bayesian generalized linear model with a zero-inflated Poisson distribution quantifying the relationship between the number of elongated follicles, diet treatment, and start date. Number of follicles ~ diet treatment + start date.

| **Model Term** | **Rhat** | **Bulk Effective Sample Size** | **Tail Effective Sample Size** |
|---|---|---|---|
| Intercept (high diet treatment, start date 1) | 1.00 | 7597 | 7103 |
| Medium diet | 1.00 | 7554 | 7612 |
| Low diet | 1.00 | 6982 | 7161 |
| Extra low diet | 1.00 | 7160 | 6695 |
| Start date 2 | 1.00 | 7319 | 7070 |
| Start date 3 | 1.00 | 7387 | 7264 |
| Start date 4 | 1.00 | 7327 | 7121 |

Table S12: Summary of mean number of elongated follicles, standard deviation, and total number of females examined per diet treatment, per start date. Ovaries of females from our anautogenous population fed the high diet treatment were also examined and confirmed no development (n = 20).

| **Diet Treatment** | **Start Date** | **Mean** | **Std. Dev.** | **Number** |
|---|---|---|---|---|
| Extra Low | 1 | 21.4 | 10.64 | 5 |
| | 2 | 13 | 8.23 | 13 |
| | 3 | 17.47 | 8.22 | 17 |
| | 4 | 9.75 | 9.79 | 16 |
| | **All Dates** | **14.29** | **9.55** | **51** |
| Low | 1 | 27.88 | 12.31 | 17 |
| | 2 | 32 | 6.67 | 13 |
| | 3 | 30.36 | 9.43 | 22 |
| | 4 | 23.06 | 9.82 | 18 |
| | **All Dates** | **28.19** | **10.24** | **70** |
| Medium | 1 | 42.24 | 21.24 | 17 |
| | 2 | 37.65 | 12.21 | 17 |
| | 3 | 41.28 | 8.14 | 18 |
| | 4 | 36.3 | 14.13 | 10 |
| | **All Dates** | **39.74** | **14.53** | **62** |
| High | 1 | 59.08 | 14.49 | 12 |
| | 2 | 41.73 | 18.46 | 15 |
| | 3 | 47 | 11.28 | 19 |
| | 4 | 45 | 7.61 | 16 |
| | **All Dates** | **47.55** | **14.29** | **62** |

Table S13: Estimates of beta coefficients from Bayesian generalized linear model with a binomial distribution quantifying the relationship between failure to produce elongated follicles and diet treatment. Number of females without developed follicles | Number of trials (with + without developed follicles) ~ diet treatment. Overlap of credible intervals with 0 indicates the beta coefficients (effect sizes) are not different from zero.

| **Model Term** | **Estimate of beta coefficient** | **Estimated error** | **Lower 95% credible interval** | **Upper 95% credible interval** |
|---|---|---|---|---|
| Intercept (high diet treatment) | -4.53 | 1.21 | -7.43 | -2.72 |
| Medium diet | 0.95 | 1.45 | -1.69 | 4.23 |
| Low diet | 0.84 | 1.43 | -1.68 | 4.03 |
| Extra low diet | 3.10 | 1.26 | 1.14 | 6.09 |

Table S14: Rhat and effective sample sizes for Bayesian generalized linear model with binomial distribution quantifying the relationship between failure to produce elongated follicles and diet treatment. Number of females without developed follicles | Number of trials (with + without developed follicles) ~ diet treatment.

| **Model Term** | **Rhat** | **Bulk Effective Sample Size** | **Tail Effective Sample Size** |
|---|---|---|---|
| Intercept (high diet treatment) | 1.00 | 3674 | 3486 |
| Medium diet | 1.00 | 3794 | 3723 |
| Low diet | 1.00 | 3855 | 4122 |
| Extra low diet | 1.00 | 3764 | 3494 |

Table S15: Estimates of beta coefficients from Bayesian generalized linear models with a gaussian distribution quantifying the relationship between the number of elongated follicles and larval development time for the low and extra-low diet treatments. Separate models were generated for each diet treatment: Number of follicles ~ Larval development time. Overlap of credible intervals with 0 indicates the beta coefficients (effect sizes) are not different from zero.

| **Diet** | **Model Term** | **Estimate of beta coefficient** | **Estimated error** | **Lower 95% credible interval** | **Upper 95% credible interval** |
|---|---|---|---|---|---|
| Low (Model1) | Intercept | 22.43 | 10.18 | 2.50 | 42.29 |
| Low | Larval Dev Time | 0.58 | 1.02 | -1.41 | 2.57 |
| XLow (Model2) | Intercept | 3.96 | 10.55 | -16.53 | 24.53 |
| XLow | Larval Dev Time | 0.70 | 0.68 | -0.62 | 2.01 |

Table S16: Rhat and effective sample sizes for Bayesian generalized linear models with a gaussian distribution quantifying the relationship between the number of elongated follicles and larval development time for the low and extra-low diet treatments. Separate models were generated for each diet treatment: Number of follicles ~ Larval development time.

| **Diet** | **Model Term** | **Rhat** | **Bulk Effective Sample Size** | **Tail Effective Sample Size** |
|---|---|---|---|---|
| Low (Model1) | Intercept | 1.00 | 10946 | 10840 |
| Low | Larval Dev Time | 1.00 | 11018 | 10943 |
| XLow (Model2) | Intercept | 1.00 | 11356 | 10406 |
| XLow | Larval Dev Time | 1.00 | 11362 | 10374 |

Table S17: Gene IDs and qPCR primer sequences

| Gene | sequence source | accession # or ID | f qpcr primer | r qpcr primer | notes |
|---|---|---|---|---|---|
| *EF1a* | Vectorbase | CPIJ009303 | TGATTGGACACGTCGATTCC | CATCTCCTGGGCTTCCTTCT | ref gene used for normalization |
| *RPL32* | Vectorbase | CPIJ001220 | TGACAAACTTGCACCAAACTG | CGTTGTGCACCAGGAACTTC | candidate ref- used in normfinder assay but not selected |
| *alphatub 84B* | Vectorbase | CPIJ019940 | ACTGCCTGGAGCATGGAATC | CTCATCCACCACGGTCGGTT | candidate ref- used in normfinder assay but not selected |
| *RPS18* | Vectorbase | CPIJ014165 | ACGAGGAGGTCGAGAAGATC | CAGTCGCTCCAGATCCTCAC | candidate ref- used in normfinder assay but not selected |
| *RPL19* | Vectorbase | CPIJ014540 | AGTTCCCTCAAGCTCCAGAAG | CTTGATCAGCTTGCGGATGCT | candidate ref- used in normfinder assay but not selected |
| *appl* | Vectorbase | CPIJ014395 | AGGAAGCGGAACCGAAGATG | CGAAGGCCAGCGTAAAGTAC | candidate ref- used in normfinder assay but not selected |
| *ilp1* | GenBank | FJ266014 | TCGCCGTGGAGGAAGAAGTC | CTTGATTTCGGCGAGCAACTG | primers for qpcr gene expression assays |
| *ilp2* | Vectorbase | CPIJ018049 | CACCATCGATCAGAGTACCTGTC | CTCGTCGTAGATGCCTTCCC | primers for qpcr gene expression assays |
| *ilp3* | NCBI | XM_001868223.1 | CGCCTCTGTGGAAGACACTTG | CAAAGTCGATCGTTGACTTCTTGAC | primers for qpcr gene expression assays |
| *ilp3* alt | GenBank | FJ266015.1 | TCAAGAAGTCGACGATCGAC | CATCATCGAGTCCACCGAGTTG | designed based on gene annotated as ILP2 (genbank), but orthologous to ILP3*. |
| *ilp4* | Vectorbase | CPIJ0108052 | CTTTCCGCAACGGTTTACAAC | CGGATAGTTCTCCGGAAACATC | primers for qpcr gene expression assays |
| *ilp5* | GenBank | FJ266016.1 | GGAGGAACAAGTGTCGGAAA | CAGCACTCCTGCGTAATGGAA | primers for qpcr gene expression assays |
| *ilp6* | NCBI | XP_001845055.1 | AGGTTTGGACCCGAACAAATCG | CACCTCGAACGGAACGGTGTT | primers for qpcr gene expression assays |
| *foxo* | NCBI | XM_001867099.1 | ACTGGGCCTCGCAACCAATC | CGTTGCCGAAAGTCAGGCGATA | primers for qpcr gene expression assays |
| *oeh* | NCBI | XP_001870999.1 | ACCCACCAATGTCCTGGAG | GCACTTGCCGTACATGATGA | primers for qpcr gene expression assays |
| *vg1b* | vectorbase | CPIJ10191 | CTACTCCTCCTCGCCTTGGT | ATGGTCCTCGACGTGACATT | primers for qpcr gene expression assays |

*After this discrepancy was detected, data saved to support ILP3 dataset

### *Supplemental Figures*

```
CLUSTAL multiple sequence alignment by MUSCLE (3.8)

"CpipILP2" ACM66967.1      ----------------LALTILAATA----------------MVQAAEQRLCGRHLVNAL
CquiILP3 XP_001868258.1   -----MNSSRSSMLALALTILAATA----------------MVHAAEQRLCGRHLVNAL
CquiILP2_XP_001868257.1   MRSASVSLSCATLVVLLLSLLGDDADASPLPNGAGASDGSDLLIHHMTRSRYCGRKLTETL
                                          * *::*.  *                : : : .* ***:*.::*

"CpipILP2" ACM66967.1      AMLCD-EFPDLHFGVKKSTIDFDRMVDYPVDDWMASMMDSSATNSVDSMMDPS--SQQLTK
CquiILP3 XP_001868258.1   AMLCD-EFPDLHFGVKKSTIDFDRMVDYPVDDWMASMMDSSATNSVDSMMDPSMQSQQQTK
CquiILP2 XP_001868257.1   AMLCQGRYP--------------MMSHHRSEYLSDQPQQPVEVEVATLPDSGVTSRG---
                          ****:  :*              *:.:  .::::.  :...  .* :: *..  *.

"CpipILP2"_ACM66967.1      MPVWMTMMYPQNYGFRAASNRNDLIPARFRKSVHSGIVDECCLRSCSINQLMKYCKTVNA
CquiILP3_XP_001868258.1    MPVWMTMMYPQNYGFRAASNRNDLIPARFRKSVHSGIVDECCLRSCSINQLMKYCKTVNA
CquiILP2_XP_001868257.1    --------YPHS-------------RSAHRRVRREGIYDECCKKSCSYTQLKSYCE----
                                   **:.              : .*.  ..** **** .*** .** .**:
```

Figure S1: Sequences for *Cpip-ilp2* (Genebank accession ACM6967.1), *Cqui-ilp2* (XP_001868257.1) and *Cqui-ilp3* (XP_001868258.1) were aligned by MUSCLE in CLUSTAL. The gene formerly named *Cpip-ilp2* is nearly identical to *Cqui-ilp3* but shows little homology to *Cqui-ilp2*, prompting a name change throughout this work.

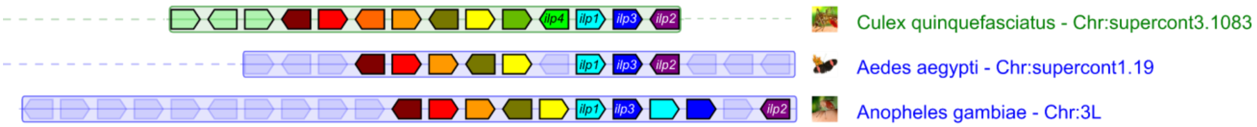


Figure S2: Synteny analysis supports an evolutionary scenario in which *Cqui-ilp4* is a lineage-specific gene duplication event. Synteny of co-linear *ilp* genes was assessed and visualized using *ilp1* as a reference gene in Genomicus Metazoa (web-code version: 2014-07-06, database version: 30.01, https://www.genomicus.biologie.ens.fr/genomicus-metazoa-30.01/cgi-bin/search.pl). Orthologous genes are indicated by matching color. *ilp* gene orthologs are labeled. Although *ilp1*, *ilp2*, and *ilp3* have clear othologs in the three mosquito species assessed with overall high conservation of nearby genes, *ilp4* is unique in *Culex.*

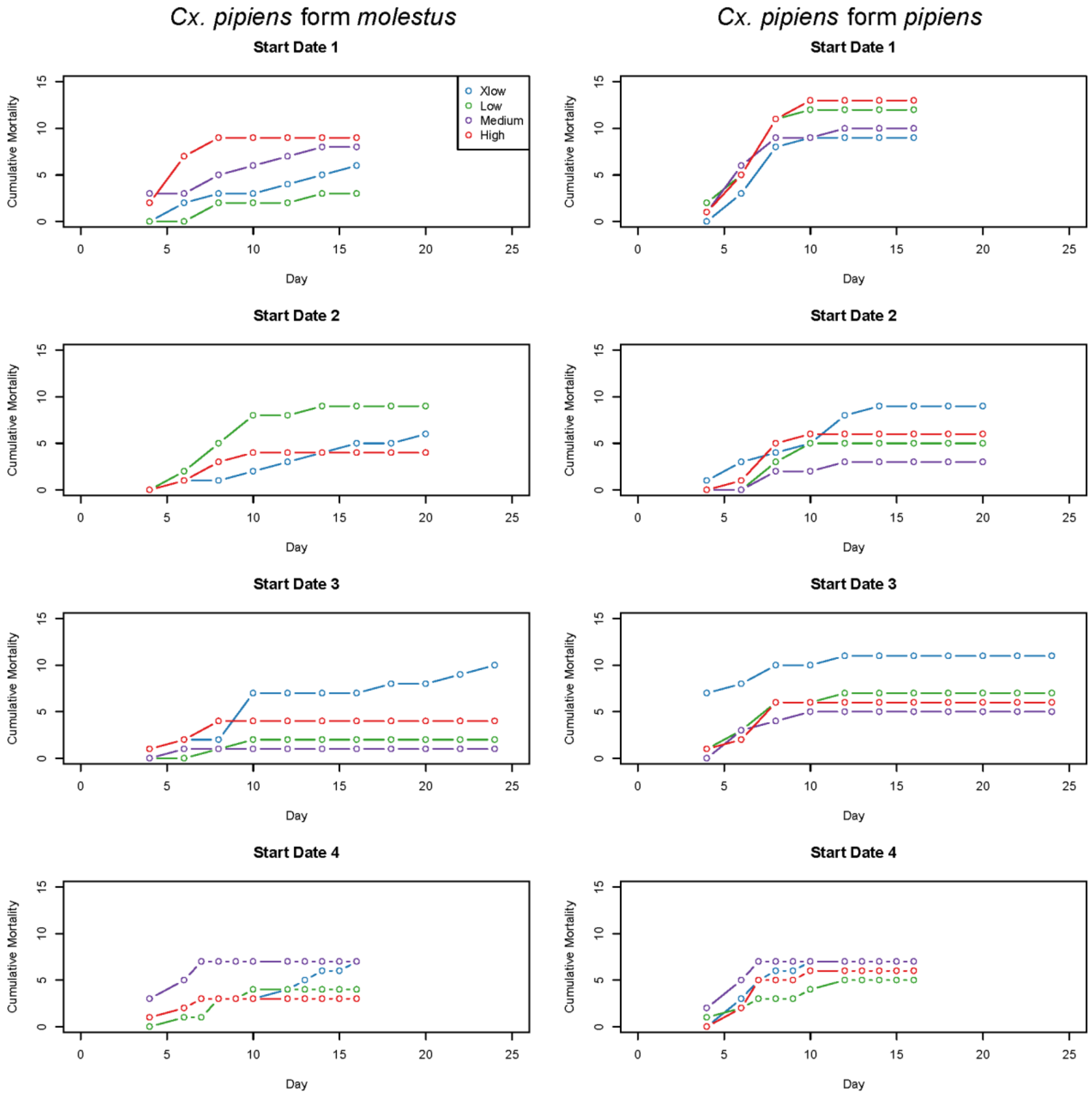


Figure S3: Cumulative larval mortality by diet treatment and start date. Panels on the left depict mortality for *Cx. pipiens* form *molestus*, while panels on the right depict mortality for *Cx. pipiens* form *pipiens*. Each replicate of the experiment concluded when the last larva pupated or died, which resulted in a different assay end point for each replicate.

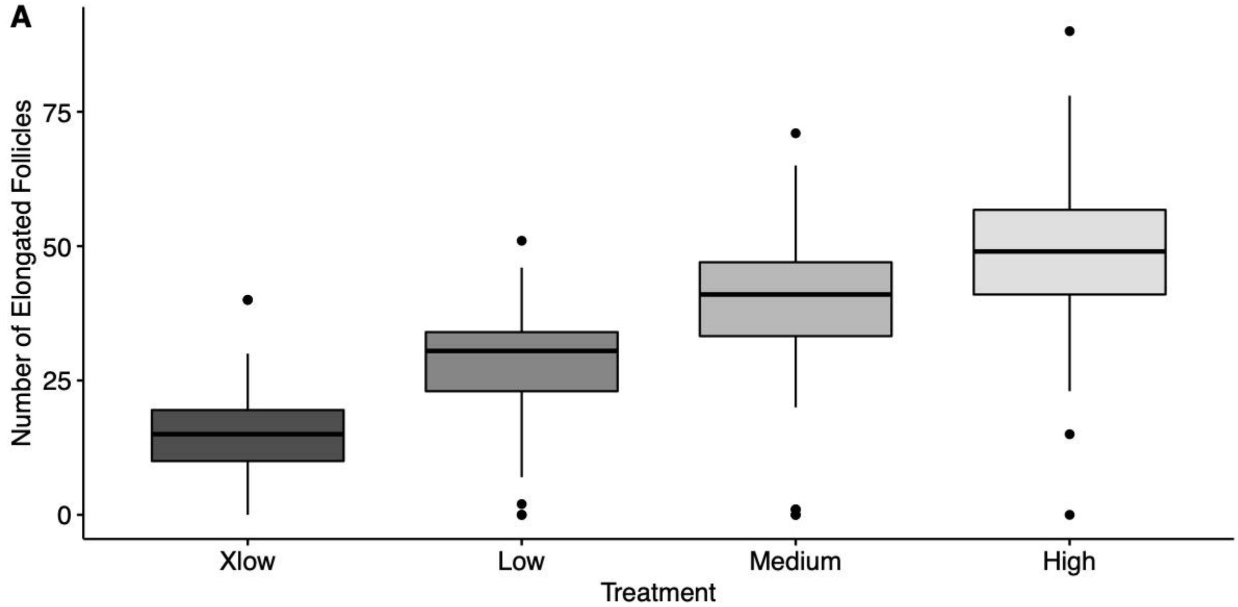


Figure S4: Larval nutrition impacts *Cx. pipiens* form *molestus* fecundity. Box plot showing the median and interquartile ranges for the number of elongated follicles per female, per treatment. Lines (whiskers) extend to 1.5 times the interquartile range, while points represent data points greater than 1.5 times the interquartile range.  Female *Cx. pipiens* form *molestus* from each treatment, and each start date, were dissected and the number of elongated follicles were counted under a dissection microscope. All females were denied an oviposition resource, and were dissected a minimum of 4 days post eclosion.

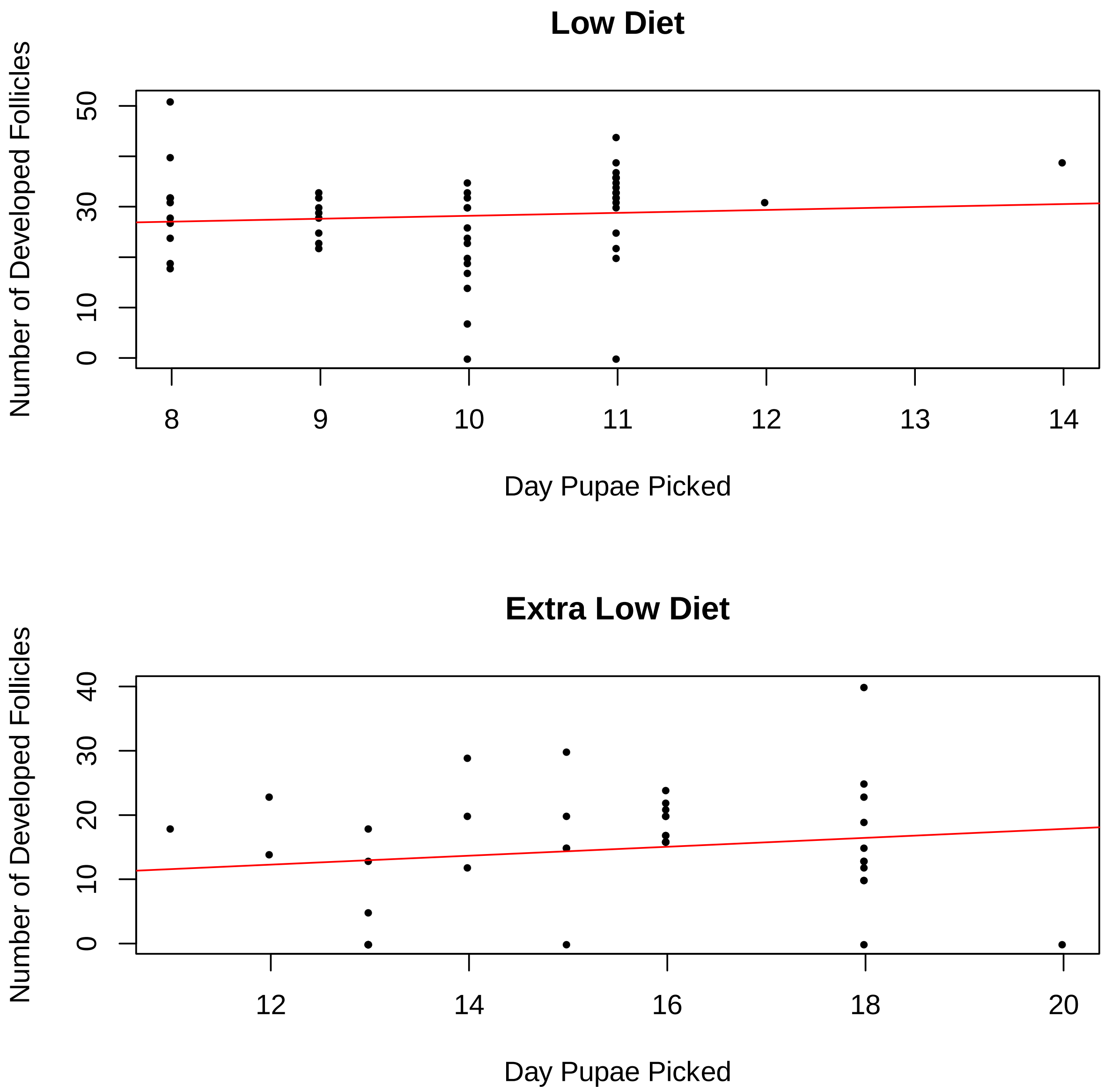


Figure S5: For *Cx. pipiens* form *molestus* females raised under low and extra low diet treatments, there was no clear relationship between time to pupation and number of elongated follicles. Points represent the number of elongated follicles according to development time (within 24 to 48 hours of pupation) for start dates 2,3, and 4. A regression line illustrating the relationship is shown red.
